# FLIGHTED: Inferring Fitness Landscapes from Noisy High-Throughput Experimental Data

**DOI:** 10.1101/2024.03.26.586797

**Authors:** Vikram Sundar, Boqiang Tu, Lindsey Guan, Kevin Esvelt

## Abstract

Machine learning (ML) for protein design requires large protein fitness datasets generated by high-throughput experiments for training, fine-tuning, and benchmarking models. However, most models do not account for experimental noise inherent in these datasets, harming model performance and changing model rankings in benchmarking studies. Here we develop FLIGHTED, a Bayesian method of accounting for uncertainty by generating probabilistic fitness landscapes from noisy high-throughput experiments. We demonstrate how FLIGHTED can improve model performance on two categories of experiments: single-step selection assays, such as phage display and SELEX, and a novel high-throughput assay called DHARMA that ties activity to base editing. We then compare the performance of standard machine-learning models on fitness landscapes generated with and without FLIGHTED. Accounting for noise significantly improves model performance, especially of CNN architectures, and changes relative rankings on numerous common benchmarks. Based on our new benchmarking with FLIGHTED, data size, not model scale, currently appears to be limiting the performance of protein fitness models, and the choice of top model architecture matters more than the protein language model embedding. Collectively, our results indicate that FLIGHTED can be applied to any high-throughput assay and any machine learning model, making it straightforward for protein designers to account for experimental noise when modeling protein fitness.

## INTRODUCTION

Machine learning (ML) approaches have been remarkably successful at solving a variety of protein design problems^1,2^. Many of these approaches involve training or fine-tuning protein design models on data generated by high-throughput experiments to optimize the desired protein function^1,3–5^. Since performance improves with training on progressively larger datasets, and biodesign models are currently thought to be more limited by the available data than by the number of parameters^6^, acquiring new datasets by measuring molecular activities in high-throughput is essential for superior protein design. High-throughput protein fitness measurements are also broadly used for benchmarking and *in silico* retrospective model performance evaluations^7–11^.

However, high-throughput laboratory experiments are inherently noisy. For example, single-step selection experiments such as phage display and other binding-based assays exhibit substantial noise in the measured enrichment ratio^12–15^; concretely, the highest-performing variants in a given single-step selection experiment show essentially 0 correlation between measured and true fitness. Despite warnings in the literature, most ML protein modeling efforts completely ignore any experimental noise present in large fitness datasets, which is tantamount to ignoring error bars. Using noisy datasets for training, benchmarking, and evaluating models can lead to inaccurate conclusions and subpar performance, as ML models can train on and potentially overfit to noise in a given dataset.

Past work on incorporating experimental noise into protein fitness modeling has primarily focused on binding-based deep mutational scanning assays. Some studies have attempted fully analytic solutions to this problem, which typically either limits the downstream modeling to linear regression^12^ or forces assumptions about the experimental parameters^13^. Other related work has focused on mitigating sequencing noise, optimizing library design, and performing multiple rounds of selection, which we do not explicitly model here^14–16^. However, the field lacks a general method for accounting for experimental noise that can be easily applied to any experimental method and ML model. We hypothesize that a generalizable framework for accounting for experimental noise could universally improve the performance of any downstream modeling on protein fitness datasets.

In this paper, we present FLIGHTED (Fitness Landscape Inference Generated by High-Throughput Experimental Data), a new approach based on Bayesian modeling to account for experimental error when generating fitness landscapes, or datasets that map a given protein sequence to fitness, from noisy high-throughput experimental data, i.e. from phage display. FLIGHTED serves as a general preprocessing step for using noisy high-throughput experimental data and is broadly applicable to any high-throughput experiment and any arbitrarily complex ML model. We apply FLIGHTED to single-step selection experiments and a new molecular recording assay and show that in both cases it generates fitness landscapes with robust, calibrated errors. We then generate two FLIGHTED-calibrated landscapes on the GB1 protein and TEV protease and benchmark a number of standard ML protein fitness models on these landscapes to learn how to improve protein fitness prediction. Our results show that using FLIGHTED improves model performance and changes relative rankings; further, data size is the most limiting factor when determining the performance of protein fitness models.

## RESULTS

### Training a FLIGHTED Model

The standard approach to machine learning for protein design takes a dataset with a single absolute fitness value corresponding to each individual protein sequence, ignoring experimental uncertainty, and uses it to generate a landscape with a single absolute fitness value for every nearby protein sequence. In contrast, FLIGHTED applies Bayesian modeling to generate probabilistic fitness landscapes in which the predicted fitness for each sequence is represented by a probability distribution (see Figure 1a). In practice, we approximate these probability distributions as normal distributions and produce means and variances for each protein sequence.

**Figure 1:**
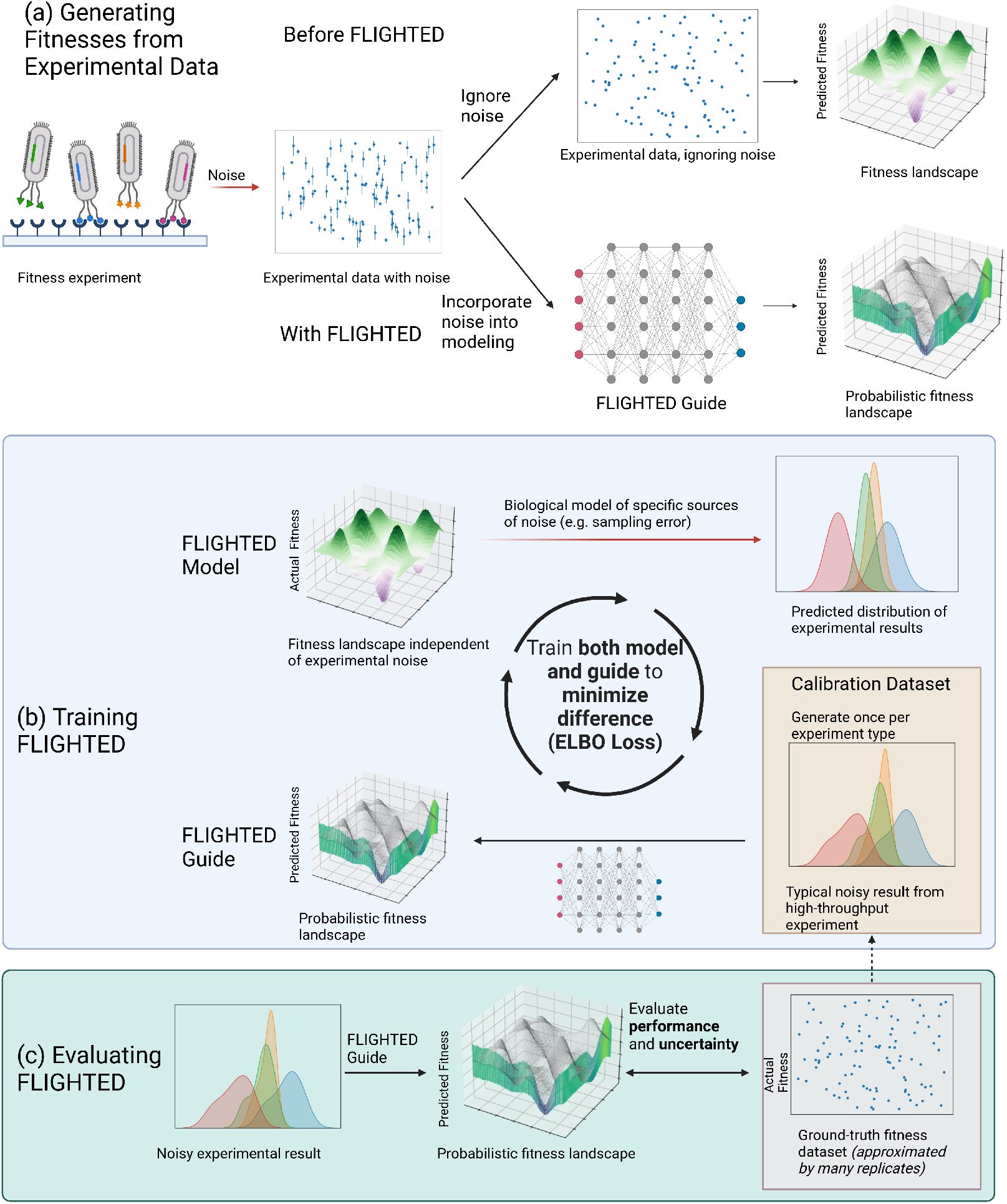
Summary of the FLIGHTED Approach to Incorporating Experimental Noise from High-Throughput Experiments. (a) A high-throughput experiment without FLIGHTED generates a readout that measures fitness and incorporates considerable noise. Before FLIGHTED, practitioners generally ignored this noise when generating a fitness landscape. With FLIGHTED, practitioners can generate a probabilistic fitness landscape that accounts for experimental noise in these high-throughput datasets. (b) To train FLIGHTED, the FLIGHTED model models a specific source of noise in a particular high-throughput experiment and generates a distribution of noisy experimental results from a fitness landscape. We then use the FLIGHTED guide (a neural network) to convert experimental readouts to fitness predictions with calibrated errors. We train both the model and the guide to minimize the difference, i.e. the ELBO loss between the two distributions of experimental results. Training uses experimental results from a calibration dataset but does not require ground-truth fitness values. (c) FLIGHTED guide is evaluated using ground-truth fitness values from a calibration dataset for predictions and uncertainty. FLIGHTED guide can then be used to measure fitnesses from any high-throughput experiment of the same type, beyond the calibration dataset.

To translate the measured activities and experimental uncertainties of sequences in the training dataset into probability distributions describing the fitness of never-measured sequences, FLIGHTED explicitly models the known sources of experimental noise for each major type of experiment. As a result, a different version of FLIGHTED must be trained for each type of experiment. All subsequent datasets generated by experiments of that type, including in other laboratories, can use that version of FLIGHTED without further modification.

Training FLIGHTED requires 1) knowledge of major sources of experimental noise for a given type of experiment, and 2) a noisy “calibration dataset” generated by running a high-throughput experiment of that type. Training yields the FLIGHTED model, which uses the our physical knowledge of noise to computationally generate a predicted distribution of experimental results from a noiseless fitness landscape, and the FLIGHTED guide, which predicts a probabilistic fitness landscape from noisy experimental results (see Figure 1b). The FLIGHTED guide and the FLIGHTED model are trained together to minimize the difference. As long as we can accurately describe the experimental noise in the calibration dataset, the FLIGHTED guide will generate a probabilistic fitness landscape that accurately accounts for the noise, which should theoretically improve accuracy over standard approaches that do not account for noise.

Evaluating FLIGHTED involves comparing the probabilistic fitness landscape generated by the FLIGHTED guide to the ground-truth fitness – which by definition is unknown, but can be approximated by averaging the results of a very large number of experimental replicates. Importantly, FLIGHTED is trained in a fully unsupervised fashion using only noisy experimental results from the calibration dataset; ground-truth fitness values are only used for evaluation. To ensure that the ground-truth fitness values are not used in training, the calibration dataset must be separated from any data used to approximate the ground-truth before training begins (see Figure 1b-c).

In this paper, we train two versions of FLIGHTED: one for single-step selection experiments, and one for DHARMA, a molecular recording assay that links fitness to base editing mutations in a canvas^17^. These versions of FLIGHTED can be used by other researchers intending to train on datasets generated by single-step selection experiments or by DHARMA.

We now describe the mathematical framework that justifies FLIGHTED; see Figure 2b. Abstractly, a high-throughput experiment generates a library of protein variants *x* that map onto fitness values *z* defined by their molecular activity under a certain set of conditions, but the noisy experimental process imperfectly generates a noisy result *y* from the actual fitness (see Figure 1, top). For each experiment, we can write down a probabilistic graphical model for this noise generation process *z → y* using our biological knowledge. That is, we generate an mathematical model for each source of noise and use it to compute a distribution of possible experimental results *y* given the fitness *z* of a protein sequence. This constitutes the FLIGHTED model for a given experiment (see Figure 1b); any sources of noise not modeled in this process will not be accounted for by FLIGHTED. For example, in the context of single-step selection experiments, we model sampling noise, or the noise that occurs when sampling from a larger population to sequence specific variants. Other sources of noise in the experiment, like proteins that bind to plastic instead of the desired molecule, are not modeled and cannot be accounted for.

**Figure 2:**
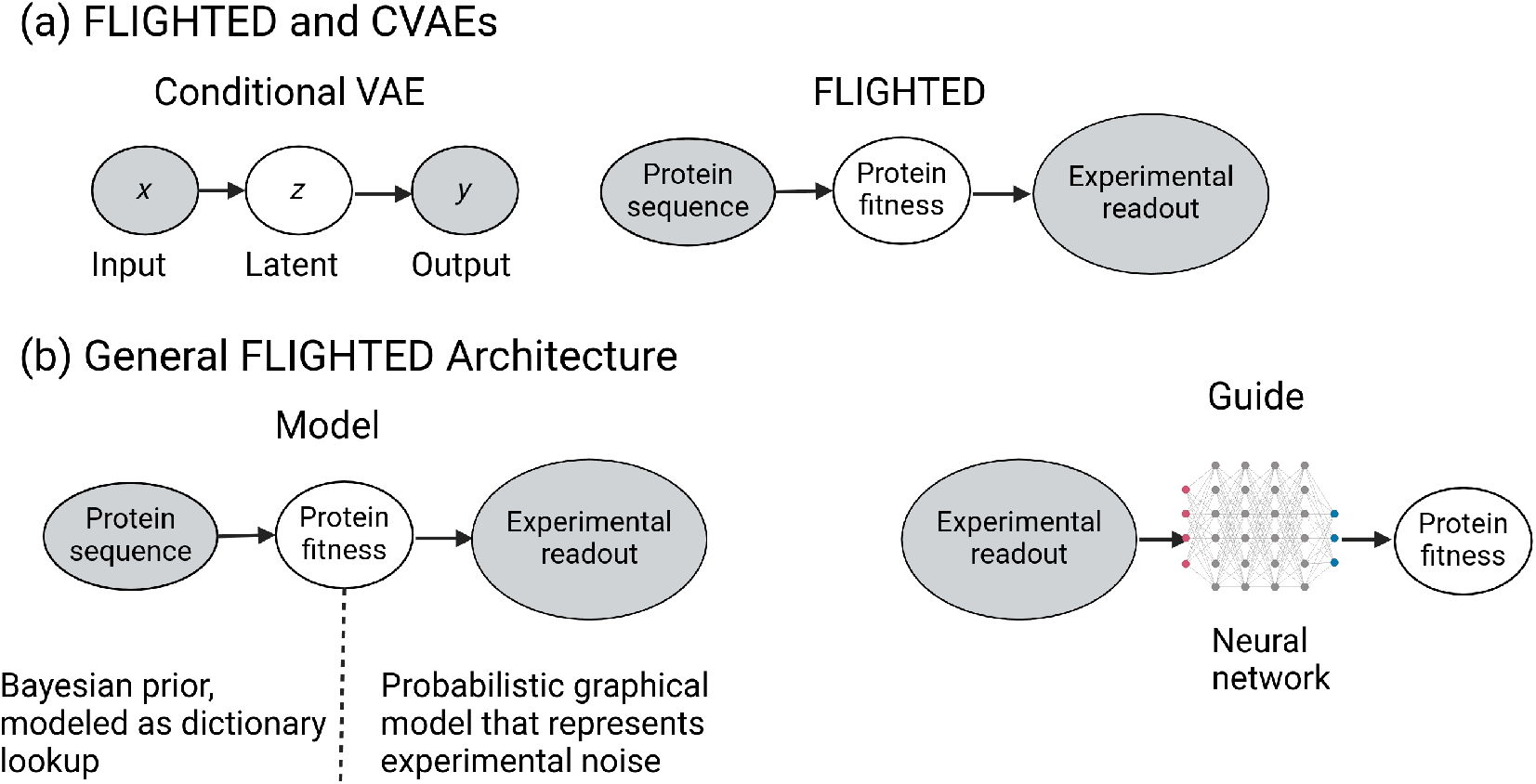
Architecture of a General FLIGHTED Model. (a) The probabilistic graphical models between FLIGHTED and the conditional variational autoencoder (CVAE) are analogous, motivating our approach, even though the applications of the two are completely different. (b) By analogy, the FLIGHTED model predicts the experimental readout from protein sequence via a sequence-to-fitness function (implemented as a dictionary lookup) and a probabilistic graphical model that models experimental noise. The FLIGHTED guide predicts protein fitness from the experimental readout using a neural network.

Because running inference on an arbitrary noise generation model to determine the fitness of each variant is computationally intractable, we use stochastic variational inference (SVI) to generate a FLIGHTED guide, or variational distribution, that predicts probabilistic fitness 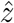 with error from experimental results *y* ^18^. Specifically, we treat the fitness *z* of each variant as a local latent variable. To avoid an arbitrarily large latent space, we use the amortization trick to learn a guide function that maps experimental results *y* to predicted fitness values 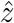 instead; this is the same trick that is used in the variational autoencoder (VAE) and the conditional variational autoencoder (CVAE)^19,20^. The guide function approximates the posterior distribution of fitness as a normal distribution. We then train our model and guide simultaneously to minimize the ELBO loss between the guide-predicted fitness values 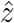 and the fitness landscape *z*, given noisy experimental results from the calibration dataset. Using SVI has two main advantages. First, model parameters can be used for any number of subsequent experimental measurements after being determined once on a calibration dataset; second, all models can have arbitrary complexity (including neural networks).

We can also view FLIGHTED as analogous to the CVAE, since both have the same graphical model (see Figure 2a)^20^. In the CVAE setup, we have an input *x*, an output *y*, and a latent variable *z*. We begin with a data-specific latent prior *p*_*θ*_(*z*|*x*) and have a probabilistic graphical model *p*_*ϕ*_(*y*|*z, x*) to generate the output. Our variational distribution or guide then predicts 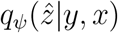. Within this framework, the ELBO loss may be written as

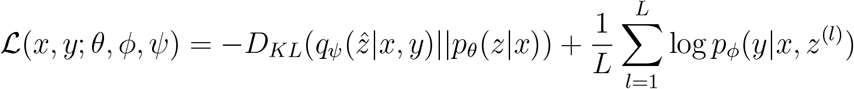

where *L* is the number of samples or particles used to approximate the loss, *z*^(*l*)^ represents a given sample of the latent variable, and *θ, ϕ, ψ* represent learnable parameters. By analogy, here the sequences are the input, the experimental result is the output, and the fitness is the latent variable. FLIGHTED’s setup is the same as the CVAE, except instead of using an arbitrary neural network to compute *p*_*ϕ*_(*y z, x*), we write down a probabilistic graphical model based on biological knowledge of the experimental process. The use of this probabilistic graphical model explains why learning from only experimental readouts – and not ground-truth fitness – is possible: our biological knowledge of the sources of experimental error constrains the space of possible models, whereas a neural network would not have any such constraints. This analogy only works because the graphical models are identical: the downstream applications of FLIGHTED and the CVAE are completely different.

To evaluate FLIGHTED’s performance (see Figure 1c), we measure both performance and uncertainty prediction of the FLIGHTED guide’s predicted fitness values and errors. The FLIGHTED guide can then be used to de-noise other experimental results generated from the same highthroughput experimental assay, even those involving different molecular libraries and targets (see Figure 1a).

The procedure we have outlined allows for the generation of trustworthy fitness measurements accounting for experimental error solely from using the FLIGHTED guide on a given experiment. While we demonstrate just two applications of FLIGHTED, it can be easily generalized to other assays; the only arbitrary step in the procedure described above is selection of neural network architecture for the FLIGHTED guide.

### FLIGHTED-Selection

We now turn to the application of FLIGHTED to single-step selection assays: experimental assays in which a library of variants is produced, a selection step occurs to separate more fit variants, and the selected population is measured; see Figure 3a^21–23^. Examples of single-step selection experiments include mRNA display, phage display, and many deep mutational scanning (DMS) studies. They comprise a substantial portion of the still-limited fitness landscapes used for machine learning^7,24^. Fitness is usually measured as the enrichment ratio, or the ratio of variant sequences sampled post-to pre-selection^21^. However, the enrichment ratio is an inherently noisy measurement due to sampling noise, the noise associated with sampling the reads to be sequenced from a larger library^21^. Sampling noise is unavoidable in a single-step selection assay and the only source of noise we choose to model in FLIGHTED^21^.

**Figure 3:**
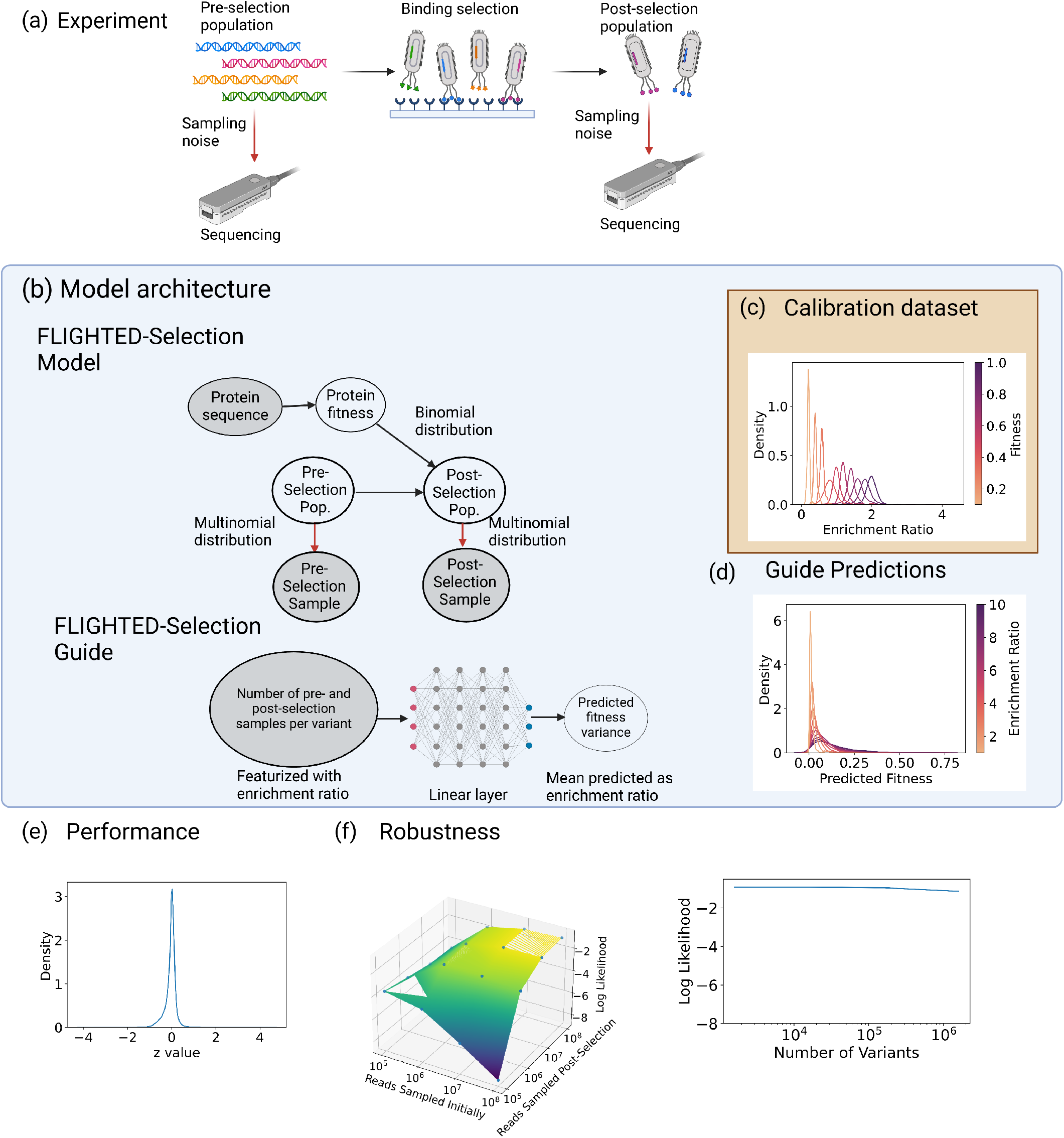
FLIGHTED-Selection Model Performance. (a) Single-step selection assays always include sampling noise when reads are sampled for sequencing. (b) The probabilistic graphical model used for FLIGHTED-Selection and model architecture for the guide. (c) Simulations indicate that the enrichment ratio has considerable noise in a single-step selection experiment due to sampling. (d) Typical FLIGHTED-Selection predictions for a given enrichment ratio. (e) FLIGHTED-Selection model predictions are well-calibrated. (f) FLIGHTED-Selection model performance is robust to changes in number of reads selected pre- and post-selection if there are more reads than variants (20^4^ here). Model performance is also robust to the number of variants.

To understand the impacts of sampling noise and generate calibration datasets for down-stream training, we ran simulations of a single-step selection assay on a randomly generated fitness landscape with 20^4^ variants. We use simulations here as a substitute of real experimental data, since we need to know the ground-truth fitness to accurately evaluate FLIGHTED-Selection’s performance; there is no easy way to measure the ground-truth fitness for any given real-world single-step selection experiment, and alternative measurement assays are generally not high-throughput and consequently fail to yield sufficient data. Figure 3c and Supplementary Figure S4a plot the probability distributions of the enrichment ratio for a given fitness, showing there is considerable noise and that this noise increases with increasing fitness.

To directly show the extent of the impact on downstream ML, we measure the Pearson *r* between fitness and enrichment ratio. Supplementary Figure S4c demonstrates that the Pearson *r* for the roughly 1000 variants with highest enrichment ratio is roughly 0, indicating single-step selection experiments offer no useful information about variants with the highest fitness. Many papers^24^ filter their data by using a minimum number of reads to reduce noise. We simulate this practice in Supplementary Figure S4c and find no substantial change. This is particularly problematic for downstream ML applications attempting to select variants with high fitness, since the most useful data points for this task are also the noisiest. Therefore, enrichment ratio is not a reliable measure of fitness.

The probabilistic graphical model for FLIGHTED-Selection is shown in Figure 3b. Given a fitness landscape and a pre-selection population, we randomly sample and sequence pre-selection, select a post-selection population based on the provided fitness values, and randomly sample and sequence post-selection. The guide model as shown in Figure 3b uses the logit of the enrichment ratio as the mean and predicts the variance using linear regression on the number of pre- and post-selection samples. FLIGHTED-Selection is trained on the calibration dataset of simulations described previously. Typical model predictions are shown in Figure 3d for given enrichment ratios; for more active variants, single-step selection experiments produce less reliable fitness measurements and FLIGHTED-Selection assigns higher variance, as expected.

To evaluate model performance, we focused on calibration, since the enrichment ratio is the optimal fitness value. We compute the *z*-value of the true fitness compared to the model prediction in Figure 3e; the resulting distribution looks very similar to a normal distribution, suggesting that FLIGHTED-Selection is well-calibrated. The mean log likelihood of our predictions is −0.97, very close to that expected for a normal distribution.

Next, we evaluated the robustness of FLIGHTED-Selection as a function of various parameters of the data generation process. In Figure 3f we showed the log likelihood as a function of number of reads sampled pre- and post-selection and demonstrated that the model was very reliable as long as the number of reads sampled was greater than the number of variants (at 20^4^). Next, we fix the number of reads sampled pre- and post-selection and evaluate robustness as a function of number of variants. In Figure 3f, we see that the log likelihood of FLIGHTED-Selection does not substantially worsen as a function of the number of variants. We also experimented with varying different parameters of the random distributions in the simulation to see whether that affected FLIGHTED-Selection model performance. In Supplementary Figure S4b, we see that these factors generally have very little effect on robustness. As a result, our use of simulated data to train FLIGHTED-Selection should not substantially affect real-world model performance.

### FLIGHTED-DHARMA

We next turn to DHARMA (Direct High-throughput Activity Recording and Measurement Assay), a recently developed high-throughput protein fitness assay^17^. DHARMA measures fitness by linking protein fitness to transcription of a base editor; this base editor is targeted to a canvas where it randomly causes C→T edits, as shown in Figure 4a. Higher fitness leads to higher base editor transcription and more C→T edits.

**Figure 4:**
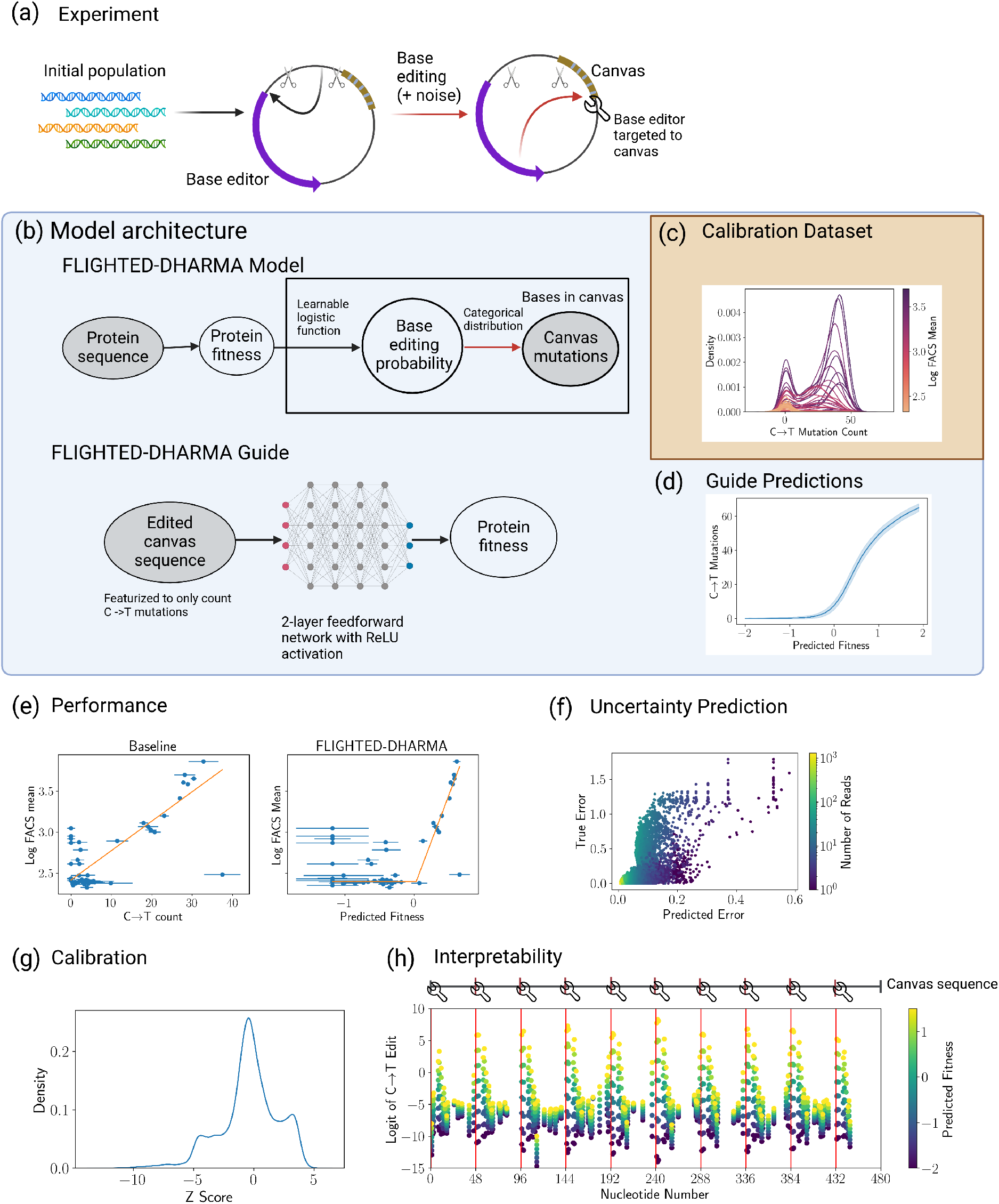
FLIGHTED-DHARMA Model Performance. (a) DHARMA links fitness to edit rates of a DNA segment (canvas) via transcriptional control of a base editor, an inherently noisy process. (b) The probabilistic graphical model for FLIGHTED-DHARMA and the architecture for the FLIGHTED-DHARMA guide. (c) Noise in DHARMA as a function of true fitness. (d) FLIGHTED-DHARMA’s prediction of number of C→T mutations as a function of fitness. (e) FLIGHTED-DHARMA improves performance of fitness predictions when compared to a baseline model that predicts number of C→T mutations. (f) FLIGHTED-DHARMA errors correlated with true error on even small subsets of DHARMA reads. (g) FLIGHTED-DHARMA is reasonably well-calibrated.(h) Canvas nucleotides most likely to be edited according to FLIGHTED-DHARMA are found at guide RNA binding sites, as expected.

DHARMA can be run in high-throughput on up to 10^6^ variants at once, since the readout is measured by cheap long-read sequencing. It can measure any protein function that can be linked to base editor activity, which includes protease activity, gene editing, and protein and DNA binding^25^. However, base editor production and activity are inherently stochastic processes^26^ so DHARMA output is noisy; Figure 4c demonstrates that repeated DHARMA experiments on the same variant can result in differing C→T edit counts. Therefore accounting for noise via an ML model like FLIGHTED is essential for reliable fitness measurements.

The probabilistic graphical model for FLIGHTED-DHARMA is shown in Figure 4b. Given a fitness landscape, we compute a base editing probability for each base in the canvas as a logistic function of fitness and then sample to determine whether each base was mutated. This model treats all canvas bases as independent but allows each base to have a different learned edit propensity. The guide predicts both the mean and variance of fitness given a single DHARMA read using a two-layer feedforward neural network. We generated a calibration dataset by running a DHARMA experiment on a 3-site library of T7 polymerase. Concretely, in Figure 4d we show the predicted number of C→T edits with error given a particular fitness.

We now assess performance and uncertainty of the trained FLIGHTED-DHARMA model. “Ground-truth” fitness measurements were produced by Fluorescence-Activated Cell Sorting (FACS) on a subset of 119 T7 variants, in which T7 polymerase drives the expression of GFP instead of base editor. Error was minimized in these FACS measurements by using a large number of cells per variant. To eliminate data leakage, FLIGHTED-DHARMA was not trained on any DHARMA read from any variant for which FACS was performed. In order to assess model performance using the FACS dataset, we must convert fitness values predicted by FLIGHTED-DHARMA to the FACS readout; since fitness is scaled arbitrarily, these two fitness values can be related by any nondecreasing function. We used a validation set to fit this function to a piecewise linear function as shown in Supplementary Figure S5a. The initial flat section of the piecewise linear functions corresponds to low-activity variants where background fluorescence dominates the FACS measurement. We then evaluated model performance on the remaining test set in Figure 4c. FLIGHTED-DHARMA was compared to a baseline consisting of the mean and variance of C→T mutation count. We found that FLIGHTED-DHARMA’s mean-squared-error (MSE) was 0.72, an improvement over the baseline MSE of 0.78. Most of the improvement is observed on the portion of the fitness landscape where both DHARMA and FACS are measuring fitness.

We also examined uncertainty by evaluating the calibration of our predicted variances. First, we directly examine calibration on the test data, using all reads available for each datapoint in the test set in Supplementary Figure S5b. FLIGHTED-DHARMA’s predicted errors increase as true error increases. The log likelihood of the baseline model on this dataset was −19.8 while the log likelihood of FLIGHTED-DHARMA was −10.0. In Figure 4d, we selected random subsets of DHARMA reads and measured the true and predicted error of FLIGHTED-DHARMA. Random subsampling tests our ability to predict error given even very small subsets of DHARMA sequencing reads. The mean log likelihood here was −3.93, suggesting that FLIGHTED-DHARMA is slightly overconfident but reasonably well-calibrated. Baseline model performance was substantially worse with a mean log likelihood of −8.00 (see Supplementary Figure S5c). Since our calibration tests included small subsets of DHARMA reads, we can be confident that FLIGHTED-DHARMA will produce reasonable errors even when given few DHARMA reads. This aids experimentalists by informing them whether a given number of DHARMA reads is sufficient for a fitness measurement.

Finally, since FLIGHTED uses a biological model of DHARMA, we can examine the parameters of that biological model directly. In Figure 4g, we compute the logit of the C→T edit probability at a given canvas residue as a function of fitness. The probability of a C→T edit increases and is more fitness-dependent every 48 residues. The base editor is targeted at these locations, so these results are expected. This is an example of how FLIGHTED can yield an interpretable model.

### Benchmarking Landscapes on FLIGHTED Data

Having established that FLIGHTED can generate fitness landscapes with reliable, calibrated errors from single-step selection and DHARMA experiments, we now turn to benchmarking ML models on the resulting landscapes. We focus on two landscapes: a 4-site landscape for the GB1 protein that measures stability and binding to IgG via an mRNA display single-step selection assay^24^, and a 4-site landscape on TEV protease that measures protease activity on the wild-type substrate using a DHARMA assay. The first landscape has frequently been used to benchmark ML models in the field^7^; we release the second landscape publicly as a source for the ML community. Basic statistics about the TEV landscape may be found in Supplementary Figures S6b - f.

The models we chose to benchmark are outlined in Figure 5a. Specifically, given a sequence, we generate an embedding from a protein language model or from a one-hot baseline^27,28^. We then feed this embedding into a task-specific top model – either a linear layer, an augmented model^29^, a feedforward neural network (FNN), a convolutional neural network (CNN), or a finetuned FNN or CNN – which is trained on this particular landscape. This strategy of blending a protein language model with a task-specific top model has repeatedly proven successful^5,7,27,28,30^. While these models are certainly not novel, to our knowledge this is the first systematic exploration of the use of CNNs as a top model.

**Figure 5:**
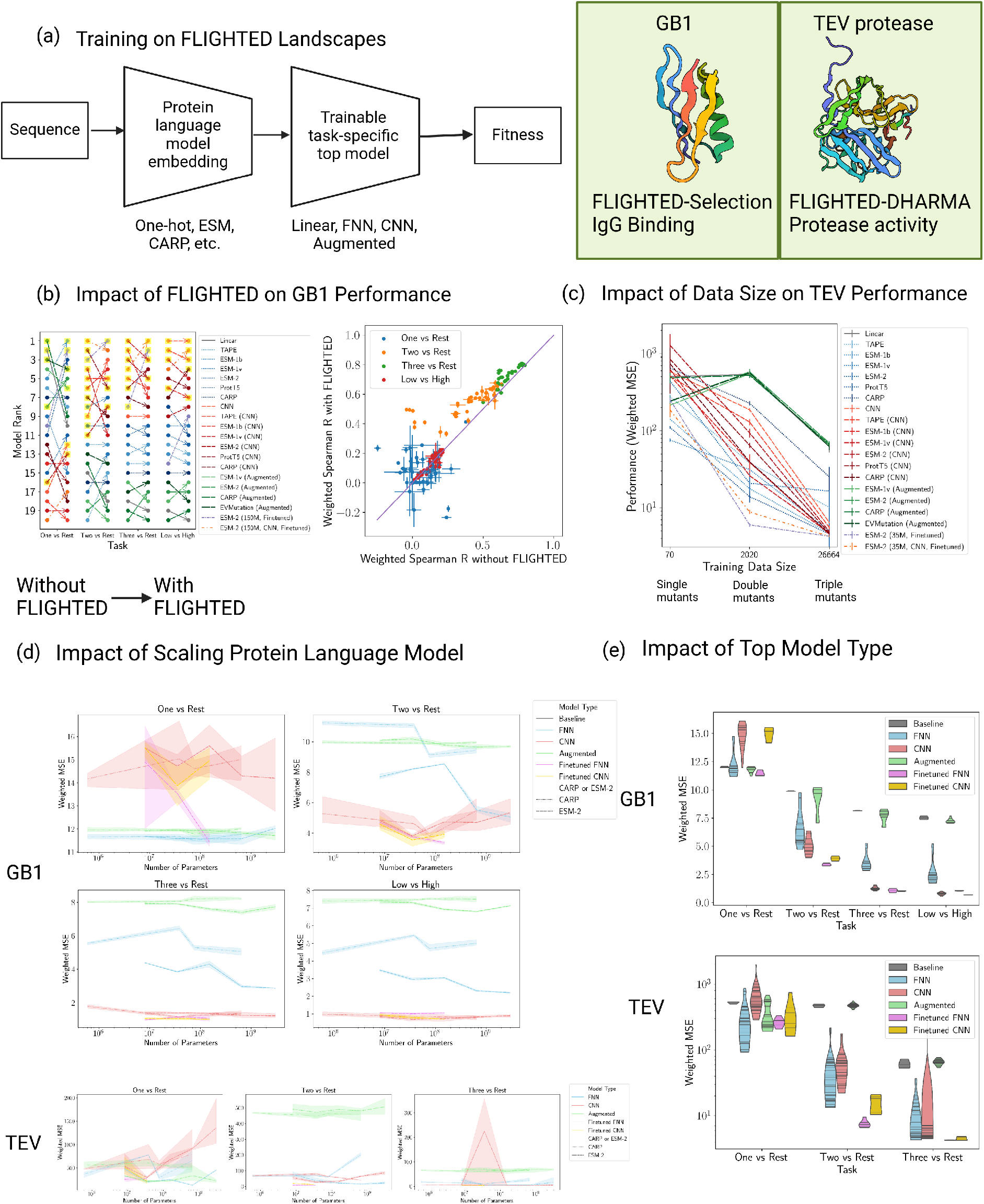
Benchmarking Protein Fitness Models on FLIGHTED Landscapes. (a) The typical model architecture we tested and the two landscapes: GB1, based on FLIGHTED-Selection^24^, and TEV protease, based on FLIGHTED-DHARMA. (b) FLIGHTED changes model ranking on the GB1 dataset, indicating the importance of de-noising when benchmarking ML models. FLIGHTED also modestly improves model performance on the GB1 dataset. (c) Exponential increases in data on the TEV dataset result in exponential increases in accuracy. (d) Scaling up protein language models does not substantially improve model performance. (e) Convolutional neural networks (CNNs) and finetuned models are the highest-performing top models across both the GB1 and TEV datasets.

To incorporate errors generated by FLIGHTED, we train and evaluate models using the inverse-variance-weighted mean squared error (MSE) as a proxy for log likelihood. We use 4 data splits on the GB1 dataset as proposed previously by the FLIP benchmark^7^: one-vs-rest, two-vs-rest, and three-vs-rest splits with only single, double, and triple mutants respectively in the training set, and a low-vs-high split with only inactive variants in the training set. We prepare similar one-vs-rest, two-vs-rest, and three-vs-rest splits of our own TEV protease dataset. See Supplementary Tables S1 and S2 for raw GB1 model performance and Table S3 for raw TEV model performance.

We first examine the impact of using FLIGHTED on models trained on the GB1 dataset in Figure 5b. The ladder plot (Figure 5b, left) shows that FLIGHTED significantly affects benchmarking outcomes. This is particularly important because of how frequently ML practitioners use single-step selection datasets to benchmark their models; without a method like FLIGHTED, the results of this benchmarking are unreliable.

Further, we also observe a general performance improvement upon using FLIGHTED data for the training set (see Figure 5b, right). Here performance is measured using weighted Spearman *ρ* so results with and without FLIGHTED can be directly compared; the weighted Spearman *ρ* accounts for variance on the test set^31^. We present correlations with FLIGHTED on the test set here; see Supplementary Figure S6a for results without FLIGHTED on the test set. 37 of the 45 models on the two-vs-rest split and 41 of the 45 models on the three-vs-rest split exhibited statistically significant improvements. We saw an average improvement of 0.15 on the two-vs-rest split and 0.05 on the three-vs-rest split. Results on the other two splits were not as consistent. These results indicate that using FLIGHTED will generally improve model performance, though this is not guaranteed.

Next, we examine the impact of various other parameters on model performance. In Figure 5c we look at the importance of data size on the TEV dataset, controlling the test set to include only quadruple mutants. As training data increases from single to double to triple mutants, model error decreases exponentially. This dramatic improvement in model accuracy indicates an increasing ability to generalize and highlights the importance of high-throughput experimental methods: with more data, our models become substantially more powerful.

We then examined which model architecture choices mattered most for performance. We found in Figure 5e that the top model was the most impactful, with all models equally unable to learn from single mutants, and CNNs performing better on larger datasets, especially three-vs-rest. Fine-tuning generally improves performance, especially for FNNs on small datasets, but is substantially more computationally expensive. Since CNNs performed well as a top model and are relatively computationally efficient compared to fine-tuning, we recommend practitioners use CNNs as a top model when they have large enough protein fitness datasets.

More surprisingly, increasing the number of parameters in the protein language model did not have a large impact on performance for most models. In Figure 5d, we ran published variants of CARP and ESM-2 with smaller numbers of parameters^28,32^. We found no substantial or consistent performance improvement, across models ranging from millions to billions of parameters, aside from fine-tuning FNNs on small datasets. Our results suggest that scaling up protein language models will not improve performance in predicting protein fitness given current data limitations. This finding has recently been corroborated^6^.

## DISCUSSION

FLIGHTED has consistently improved model performance by generating fitness landscapes with robust, calibrated errors for two very different high-throughput experiments. Since it is based on Bayesian modeling and variational inference, it can be easily extended to other high-throughput experiments or more complicated noise generation models of single-step selection by taking into account effects such as biased library design or sequencing error. We have demonstrated that FLIGHTED models can be used reliably under a wide range of experimental conditions and offer a modest performance improvement in fitness prediction. FLIGHTED results can also be used to combine fitness readouts from different experiments and optimize experimental design by maximizing signal-to-noise.

Our results on FLIGHTED-Selection suggest that common ML benchmarks like FLIP^7^ or ProteinGym^11^ need to be corrected to account for noise inherent in single-step selection. Correcting these benchmarks will change downstream evaluations of protein models and should improve performance on average.

FLIGHTED-DHARMA improves the use of DHARMA as a data generation method for machine learning. The noisy nature of biological circuits presents challenges in effectively using DHARMA to quantify activities. We provide calibrated error estimates that inform practitioners when they have enough DHARMA reads to make a reliable fitness estimate. DHARMA can be used to cheaply generate large datasets of up to 10^6^ variants using nanopore sequencing alone. Comparing FLIGHTED predictions for DHARMA and single-step selection, we see that DHARMA may be more accurate for distinguishing between highly active variants while selection experiments may be more accurate at identifying the fitness values of comparatively inactive variants.

Our benchmarking results support two important conclusions about the development of future protein fitness ML models. First, data size is currently the most important factor, underscoring the importance of gathering large, high-quality datasets through methods such as DHARMA. Second, the top model architecture matters much more than the embedding, so practitioners should focus on development of top model architecture and less on scaling up protein language models. Specifically, in the first systematic exploration of CNNs as top models, we found that they performed very well while being computationally cost-efficient. We encourage further exploration of the potential improvements that might be achieved by adopting CNNs and other poorly-studied architectures as top models.

## STAR METHODS

### Resource availability

#### Lead contact

Requests for further information and resources should be directed to and will be fulfilled by the lead contact, Kevin Esvelt.

#### Materials availability

This study did not generate new materials.

#### Data and code availability

- Model weights for FLIGHTED-Selection and FLIGHTED-DHARMA, the FLIGHTED GB1 landscape, and all GB1 models have been released at this Zenodo repository. The FLIGHTE TEV landscape and all TEV models have been released at this Zenodo repository.
- All code necessary to run FLIGHTED and reproduce all figures in this paper has been released in at this Github repository.
- Any additional information required to reanalyze the data reported in this paper is available from the lead contact upon request.

### Method Details

All models were implemented using the probabilistic programming package Pyro^33^ alongside PyTorch^34^. Use of Pyro makes model development substantially easier due to rapid iteration of model architectures.

#### Single-Step Selection

#### Dataset Generation

Single-step selection experiments were simulated on a landscape of *N*_var_ = 20^4^ variants (representing a peptide of length 4), with the *i*th variant for 1≤*i* ≤*N*_var_ having a fitness or selection probability *p*_*i*_ uniformly distributed between 0 and 1, i.e. *p*_*i*_ *∼* 𝒰([0, 1]). The total initial population *N*_init_ = 10^11^. Each variant *i* had an initial population drawn from the Dirichlet distribution *N*_init,*i*_ *∼ N*_init_Dir(*α*), *α* = (1, …, 1) with length *N*_var_. An initial pre-selection sample *N*_sampled, init,*I*_ was drawn according to a multinomial distribution Mult 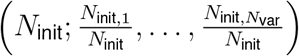, with *N*_sampled, init_ = 10^8^ samples being drawn.

In the selection step, the *i*th variant had a post-selection population drawn from the binomial distribution, i.e. *N*_final,*i*_ *∼* B(*N*_init,*i*_; *p*_*i*_) and *N*_final_ =∑_*i*_ *N*_final,*i*_. The po st-selection sample *N*_sampled, final_ was similarly drawn according to a multinomial distribution Mult 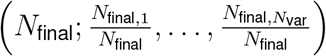 with *N*_sampled, final_ = 10^8^ samples being drawn. We ran 100 simulations on a single landscape to generate training data and to assess the noise levels in the enrichment ratio as shown in Figure 3c.

The parameters of this simulation were selected to be similar to those used in generating the GB1 dataset^24^. Simulations with smaller numbers of reads drawn both pre- and post-selection were used to assess robustness. Simulations were also run with varying Dirichlet parameters and using a beta distribution for the selection probability instead of a uniform distribution; these did not significantly affect the noise found in the enrichment ratio. Specifically, we drew *p*_*i*_ *∼*Beta(1, *b*), and varied the scalar *b* between 1 and 100. This represents fitness values that peak at lower or higher values as opposed to being uniform between 0 and 1. We also drew the initial population *N*_init,*i*_ *∼N*_init_Dir((*a*, …, *a*)) with the scalar *a* varied between 0.1 and 10, allowing for biases in the initial population towards one variant.

#### The FLIGHTED-Selection Model

The model is an implementation of the probabilistic graphical model shown in Figure 3b in Pyro. The sequence-to-fitness function is simply a dictionary lookup, with each variant *i* mapped to a fitness mean *m*_*i*_ initially sampled from 𝒩 (0, 1) and variance *σ*_*i*_ sampled from Softplus(𝒩 (0, 1)), where the softplus function is used to ensure that the variance is positive. Both *m*_*i*_ and *σ*_*i*_ are learnable. The fitness *z*_*i*_ of each variant is sampled ∼𝒩 (*m*_*i*_, *σ*_*i*_) and the probability of binding *p*_*i*_ = *σ*(*z*_*i*_), where *σ* is the sigmoid function used to ensure the probability is between 0 and 1. To avoid issues near the boundaries, fitness values are clamped to be between 0.001 and 0.999. The initial population of each variant *N*_init,*i*_ is a local parameter in Pyro. We then follow the noise generation procedure described in the first paragraph of the Data Generation Section to predict the distribution of experimental results. Note that while the noise generation procedure is mathematically identical, we are using it for different purposes; we previously used it for data generation, and are now using it for defining the probabilistic model. Robustness tests of FLIGHTED-Selection on changes to the data generation process ensure that FLIGHTED is not overly specialized to this single noise generation procedure.

The guide predicts the fitness mean for a given experiment as 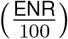, where ENR is the enrichment ratio of post-selection to pre-selection sampled reads:

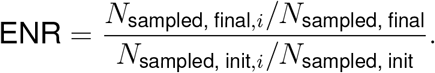

The scale factor of 100 ensures that the transformed enrichment ratio is between 0 and 1 before being fed into the logit function; it can be changed arbitrarily with no impact on model performance after retraining. The variance is predicted by a linear model with 2 inputs: 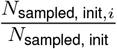 and 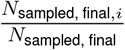. The predicted variance is also transformed by a softplus function *F* to ensure that it is positive.

Each data point for the FLIGHTED-Selection model consists of an experiment or simulation, not the outcome of a single variant within an experiment, because the number of reads of a given variant sampled pre- and post-selection depends on the population of other variants in the sample. To combine predictions from the guide model for different experiments in a single batch, we simply multiply the relevant Gaussians and sample the fitness from the resulting combined Gaussian distribution.

We used a learning rate of 10^−2^, a learning rate on the landscape model (the sequence-to-fitness function) of 10^−1^, a batch size of 10, 150 epochs, 10 particles in the ELBO loss function in Pyro, and a plateau learning rate scheduler with a patience of 4 epochs. Other hyperparameters can be found in the released code and in Table 1. The model was trained on 80 of the 100 simulations, validated on another 10 (randomly split), and tested on the final 10.

**Table 1:** Hyperparameters used for FLIGHTED-Selection.

| Hyperparameter | Value |
| --- | --- |
| Number of amino acids (alphabet length) | 20 |
| Sequence length for dataset | 4 |
| Learning rate | $10^{-2}$ |
| Sequence-to-fitness function learning rate | $10^{-1}$ |
| Batch size | 10 |
| Number of epochs | 150 |
| Test set batch size | 1 |
| Number of ELBO particles | 10 |
| Total population | $10^{11}$ |
| Sampled population | $10^8$ |
| Learning rate scheduler | Plateau scheduler |
| Plateau patience | 4 |

### DHARMA

#### T7 Polymerase Dataset Generation

We engineered a biological circuit to associate the enzymatic characteristics of T7 RNA polymerase variants with mutations accumulating on a designated DNA sequence, known as the canvas. T7 RNA polymerase is an enzyme responsible for transcribing DNA into RNA, and we constructed a library of variants with different recognition and activity profiles.

The transcription of the base editor, an enzyme that can induce mutations, is controlled by a T3 promoter. This promoter is selectively recognized by a subset of the T7 RNA polymerase variants. To moderate the expression levels of the T7 polymerase library and prevent rapid saturation of the canvas—which would compromise the collection of meaningful activity data—we used both a weak constitutive promoter and a weak ribosomal binding site.

We incorporated this biological circuit into cells already containing plasmids that express the single guide RNA (sgRNA), through a technique called electroporation. The sgRNA serves to guide the base editor to the specific canvas sequence where mutations are to be introduced. After electroporation, the cells were grown continuously in a bioreactor. Following this growth period, the region of the T7 polymerase library and the canvas with mutations was selectively amplified as contiguous fragments of DNA. The amplified material was then subjected to longread Nanopore sequencing for detailed analysis of the mutations and activities of different T7 RNA polymerase variants.

To facilitate data analysis, we implemented a data processing pipeline. Its core function is to identify, tabulate, and assign mutations present in each sequencing read to the corresponding library member. This assignment is based on internal barcodes represented as degenerate codons in the T7 RNA polymerase sequence. The pipeline accepts raw sequence reads in standard genomic formats and performs length-based filtering and optional sequence trimming. An algorithm incorporating local sequence alignment is used for barcode recognition and classification during the demultiplexing of reads. Additionally, each read is aligned to the reference sequence of the canvas to identify the location of each C→T mutation, which is then stored in a matrix for downstream ML model training.

#### The FLIGHTED-DHARMA Model

The model is an implementation of the probabilistic graphical model shown in Figure 4b in Pyro. The sequence-to-fitness function is simply a dictionary lookup, with each variant *i* mapped to a fitness mean *m*_*i*_ initially sampled from 𝒩(0, 1) and variance *σ*_*i*_ sampled from Softplus(𝒩(0, 1)), where the softplus function is used to ensure that the variance is positive. Both *m*_*i*_ and *σ*_*i*_ are learnable. The fitness *z*_*i*_ of each variant is sampled *∼ 𝒩*(*m*_*i*_, *σ*_*i*_) and fitness values are clamped at −2.

For each position *j* in the canvas, we set a learnable parameter *r*_*j*_ for a baseline rate of generic mutations (which accounts for most sequencing error from the long-read sequencing). For positions that are cytosines, we set a learnable parameter *m*_*j*_ for the fitness-dependent slope and *b*_*j*_ for the intercept. Then for all cytosines, given a fitness *z* of the variant, the logit of the C→T edit is set to *m*_*j*_ + *b*_*j*_*z* and the logit of any other mutation or deletion is *r*_*j*_. For non-cytosine residues, we simply have a logit of any mutation set to *r*_*j*_. We sample from the one-hot categorical distribution for each residue independently to get the output DHARMA read, i.e. from Cat((1, *m*_*j*_ + *b*_*j*_*z, r*_*j*_)) for cytosine residues and Cat((1,−∞, *r*_*j*_)) for non-cytosine residues.

The guide predicts fitness mean and variance from a single DHARMA read. We used a feedforward network with 2 layers with a hidden dimension of 10 and a ReLU activation between the two layers. To simplify the learning problem, only cytosine residues were fed into the guide and each position was featurized with a one-hot encoding of two classes: a C→T edit or a non-C→T edit (including both no mutation or other mutations or deletions). Variances were transformed under the softplus function. Fitness values of variants with multiple reads can be predicted by multiplying the appropriate predicted Gaussians of the individual read.

The ELBO loss used 1 ELBO particle. The landscape model (sequence-to-fitness function) learning rate was 10^−2^, the learning rate elsewhere was 10^−4^, and the batch size was 2. We used a plateau learning rate scheduler with a patience of 1 epoch and a factor of 0.1, stepped based on the training loss (not validation loss). The model was trained for 25 epochs. Other hyperparameters are found in Table 2.

**Table 2:** Hyperparameters used for FLIGHTED-DHARMA.

| Hyperparameter | Value |
| --- | --- |
| Learning rate | $10^{-4}$ |
| Sequence-to-fitness function learning rate | $10^{-2}$ |
| Batch size | 2 |
| Number of epochs | 25 |
| Test set batch size | 10 |
| Number of ELBO particles | 1 |
| Number of layers in guide | 2 |
| Guide hidden dimension | 10 |
| Guide activation | RELU |
| Learning rate scheduler | Plateau scheduler, on training loss |
| Plateau patience | 1 |
| Plateau learning rate factor | 0.1 |

To minimize data leakage, all reads corresponding to sequences in the fluorescence dataset were eliminated from training. 10% of the remaining reads were held out as a validation set to pick the optimal epoch for the model.

Many of the hyperparameters were carefully tuned using a grid search. For tuning, we wanted to use the MSE with the FACS data, so we needed to fit the predicted fitness values to the FACS data. We did not use Spearman correlation so we could measure true error (as opposed to rank-order error), and consequently we could account for situations where the function relating the FACS data to predicted fitness was not increasing (as happened here). For this, we split out a 50% of the FACS data to use as a fitness regression/validation set and evaluated the mean of the logarithm of all FACS samples. Each model was given the option of fitting using either a piecewise linear function or a linear regression. The linear regression was fit normally. The piecewise linear function corresponded to the functional form

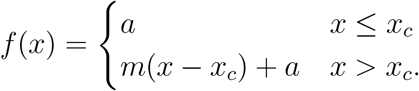

This ensured that for low fitness values, the predicted fluorescence was constant, as would be expected for background fluorescence. We then used orthogonal distance regression, as implemented in scipy, to fit the piecewise linear function accounting for errors in both predicted fitness and the FACS data.^35^ Between the linear regression and piecewise linear fit, the function with the lower MSE on the validation data was selected.

This validation set MSE was also used for selecting hyperparameters, leaving the remaining 50% of the FACS data as a test set used solely for evaluating model performance as shown in Figure 4c. Hyperparameters tuned included the batch size, learning rate, learning rate scheduler step, and number of layers in the guide model. We also evaluated the optimal number of hours to grow the cells in the bioreactor, as an example of an experimental parameter than can be tuned using FLIGHTED.

#### Model Evaluation

Model performance was measured with the 50% held-out test set of the FACS data, as described above, using the fitness values predicted by the guide model in FLIGHTED DHARMA and the fitness-to-fluorescence function fit on the validation set. The results of that performance evaluation are shown in Figure 4e.

We then evaluated calibration of the model. To increase the number of data points (since we only had 119 variants measured through FACS), we subsampled subsets of reads for each variant. Specifically, for all test set variants, we sampled 10 subsets each of sizes ranging from 1 to the maximum possible number of reads. We then predicted the fitness of each variant using the given subset of reads with the guide model. We computed the true fitness using the FACS data, eliminating any data points where the true fitness was below the baseline of the piecewise linear function. We then computed the *z*-score by comparing this predicted and true fitness. This generates the plots shown in Figure 4f and 4g.

### Benchmarking Models on FLIGHTED Datasets

#### The GB1 Dataset

To compute the corrected GB1 landscape with FLIGHTED-Selection, we took the guide and ran inference on the released GB1 landscape data, which provided both pre- and post-selection read counts in addition to the enrichment ratio^24^. We omitted all data points that were omitted in the original study due to not being observed. We then followed the published data splits provided by FLIP^7^ to generate datasets both with and without FLIGHTED-Selection for the GB1 problem. Models trained on datasets with corrections from FLIGHTED-Selection are trained with an MSE loss weighted by the inverse variance, i.e.

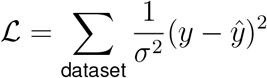

where *y* is the true value, *ŷ* is the prediction, and *σ*^2^ is the variance. Weighted MSE is used to account for the variance, assuming that the likelihood of the data is the same as that of a normal distribution with the provided variance. Performance results are regular MSEs for the datasets without corrections and weighted MSEs for datasets with corrections. Weighted Spearman *ρ* measurements were made with 100 Monte Carlo replicates^31^.

#### The TEV Dataset

High-throughput quantification of TEV protease fitness using DHARMA was made possible by a biological circuit that couples the activity of TEV protease to base editor transcription driven by a T7 promoter. As part of the circuit, the T7 RNA polymerase, normally repressed by its natural inhibitor T7 lysozyme, becomes functional after the inhibitor is disabled by active TEV protease variants, thus allowing the transcription of base editor to proceed, which in turn introduce mutations to the canvas. We engineered a variant of T7 lysozyme in which the TEV protease substrate sequence is inserted in the middle of the coding sequence. This modified T7 lysozyme, expressed on a separate plasmid under the control of a medium constitutive promoter, together with the T7 lysozyme tethered to the T7 RNAP, provides an enhanced dynamic range of base editing coupled to the activity of TEV protease variants.

The library of TEV protease used in this study was obtained by performing site-saturation mutagenesis on 4 amino acid residues (T146, D148, H167, and S170) in the TEV protease S1 pocket, which is known to interact with P1 residue on the substrate and determine substrate specificity. Briefly, NNK degenerate primers was used to introduce mutations at the targeted residues in a PCR reaction. The pool of amplicons was then cloned into a Golden Gate vector comprising the rest of the TEV protease expression cassette, the sgRNA and the canvas sequence. Commercial electrocompetent cells were then transformed with the cloning reaction, selected with appropriate antibiotics on agar plate overnight and subjected to DNA extraction. The resulting library was then sequenced to assess for bias and coverage. After quality control, the TEV protease library was transformed into electrocompetent cells that already express the base editor, the T7 RNAP and the engineered T7 lysozyme. The transformants were then selected with appropriate animate and grown continuously in a bioreactor. Multiple time points were taken during this growth period to find the optimal incubation time. The region of the plasmid containing TEV protease library and the canvas sequence was selectively amplified as contiguous fragments of DNA. To minimize the amplicon size, nucleotides not of interest were removed via self-circularization and re-amplification. The final amplified material was then subjected to longread nanopore sequencing to simultaneously retrieve both variant identity and the mutations on the canvas sequence for each individual library member. Sequencing data processing and analysis was performed as described in the T7 dataset generation section.

The TEV landscape was generated by processing the raw TEV data with FLIGHTED-DHARMA as trained on the T7 dataset. We split the dataset by mutation distance from the wild-type to create a one-vs-rest, two-vs-rest, and three-vs-rest data split. We then split out a random 10% of the training data to use as validation data. Models trained on the TEV dataset are trained with a weighted mean-squared-error (MSE) loss, weighted by the inverse variance, as above. Performance results are also weighted MSEs. In Figure 5c, we use a separate test set consisting of only quadruple mutants (i.e. the test set in the three-vs-rest split) for all models.

#### Modeling the GB1 and TEV Datasets

Our models are inspired by but go beyond the models proposed in FLIP (Fitness Landscape Inference for Proteins)^7^. See Table 3 for a list of hyperparameters. The linear regression model (labeled Linear) takes one-hot embeddings of the full sequence and runs them through 1 linear layer. It uses an Adam optimizer with a learning rate of 10^−2^, a batch size of 256, and a weight decay of 1. The CNN model (labeled CNN) takes one-hot embeddings of the full sequence and has 1 convolutional layer with 1024 channels and filter size 5, and same padding. It then has a 1D batch normalization layer and a ReLU activation, followed by an embedding neural network consisting of a linear layer to 2048 dimensions and a ReLU activation. Then there is a max-pooling layer over residues, a dropout layer with probability 0.2, and a final linear layer for the output. The CNN is trained with a batch size of 256 and an Adam optimizer with a learning rate of 10^−3^ and weight decay of 0 for the convolutional layer, a learning rate of 5*10^−5^ and weight decay of 0.05 for the embedding layer, and a learning rate of 5*10^−6^ and weight decay of 0.05 for the output layer. Unlike the original FLIP paper, we did not use early stopping and trained all models for a full 500 epochs, using validation set performance to select the optimal model.

**Table 3:**
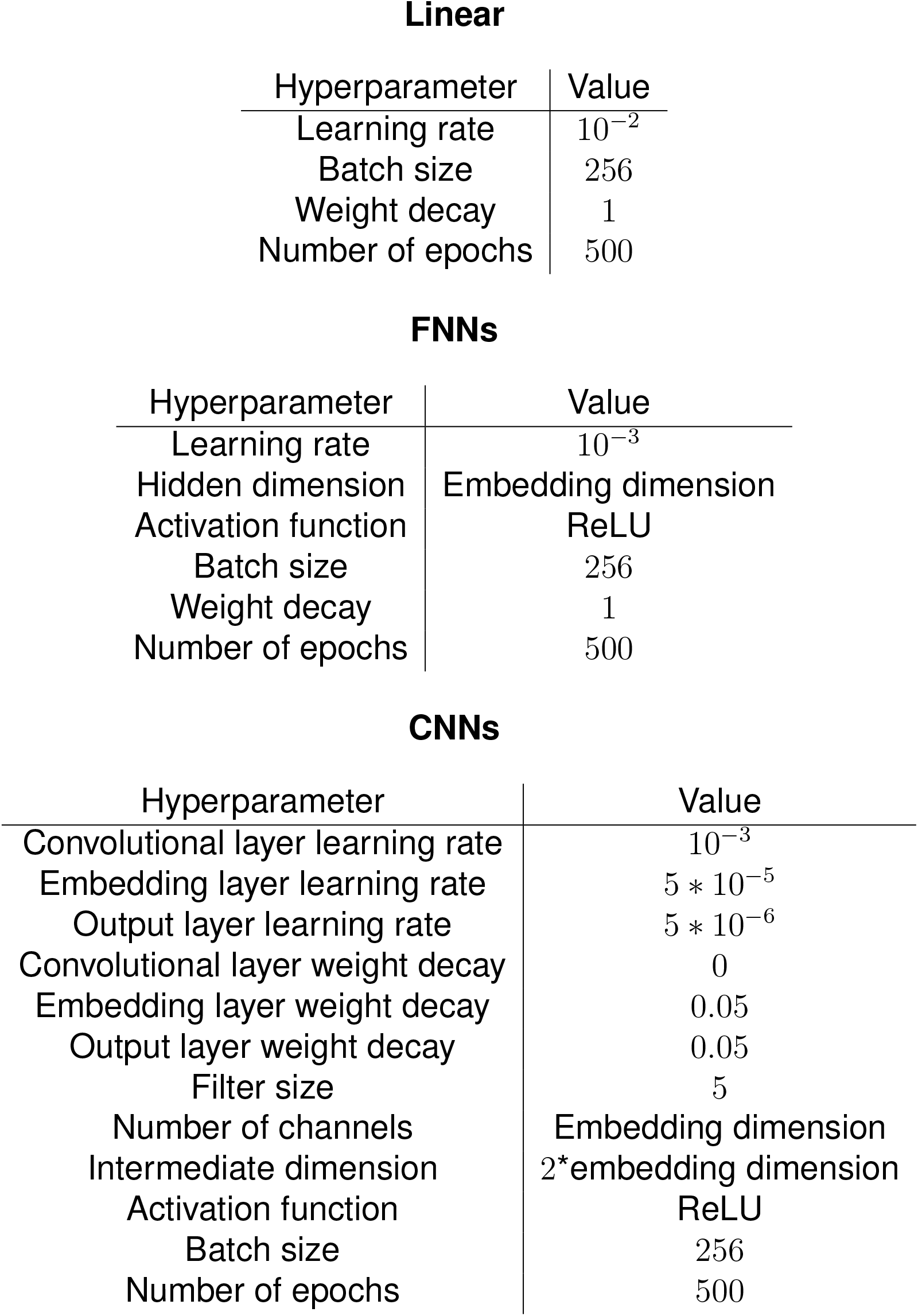
Hyperparameters used for benchmarking models on fitness datasets.

Linear
| Hyperparameter | Value |
| --- | --- |
| Learning rate | $10^{-2}$ |
| Batch size | 256 |
| Weight decay | 1 |
| Number of epochs | 500 |

FNNs
| Hyperparameter | Value |
| --- | --- |
| Learning rate | $10^{-3}$ |
| Hidden dimension | Embedding dimension |
| Activation function | ReLU |
| Batch size | 256 |
| Weight decay | 1 |
| Number of epochs | 500 |

| Hyperparameter | Value |
| --- | --- |
| Convolutional layer learning rate | $10^{-3}$ |
| Embedding layer learning rate | $5 * 10^{-5}$ |
| Output layer learning rate | $5 * 10^{-6}$ |
| Convolutional layer weight decay | 0 |
| Embedding layer weight decay | 0.05 |
| Output layer weight decay | 0.05 |
| Filter size | 5 |
| Number of channels | Embedding dimension |
| Intermediate dimension | 2*embedding dimension |
| Activation function | ReLU |
| Batch size | 256 |
| Number of epochs | 500 |

The models labeled TAPE, ESM-1b, ESM-1v, ESM-2, ProtT5, and CARP all use mean embeddings across the entire sequence that are fed into a feedforward neural network to compute the output^8,27,28,36,37^. The output feedforward neural network has 2 layers with a hidden dimension equal to the embedding dimension of the protein language model and ReLU activation. All models are trained with an Adam optimizer with batch size 256 and learning rate 10^−3^. The TAPE model used was the transformer, the ESM-1v model used was version 1, the largest ESM-2 model used was the 3 billion parameter version (due to memory issues with the larger model), the ProtT5 model was the suggested ProtT5-XL-UniRef50 model, and the CARP model was the largest 640 million parameter version. For Figure 5d, we ran smaller versions of CARP and the ESM-2 models, as specified by their parameter number, with the same architectures as described above^28,32^.

The models labeled TAPE (CNN), ESM-1b (CNN), ESM-1v (CNN), ESM-2 (CNN), ProtT5 (CNN), and CARP (CNN) use residue-level embeddings that are fed into a CNN, similar to the baseline CNN described above. The intermediate dimension output by the embedding neural network is set to be twice the dimension of the output of the protein language model. All other parameters remained the same.

The models labeled ESM-1v (Augmented), ESM-2 (Augmented), CARP (Augmented), and EVMutation (Augmented) used the zero-shot variant effect prediction from the model in question, combined with a one-hot encoding of the entire sequence that was fed into a linear layer^27,28,32,38^. These were trained with the same parameters as the baseline linear model. The ESM models used the masked marginal approach proposed in^27^ to compute zero-shot variant effect prediction. EVMutation used the default parameters^38^.

Fine-tuned models are trained as described above, but we also fine-tuned the underlying ESM model that generated the embeddings with a learning rate of 10^−6^. Due to compute and cost limitations, fine-tuning was only done for the ESM models with 8 and 35 million parameters for the TEV dataset and 8, 35, and 150 million parameters for the GB1 dataset.

We observed high run-to-run variance in the performance of many models. As a result, all models were run in triplicate and we have reported both the mean and standard deviation of model performance.

## Acknowledgments

V.S. acknowledges funding from the Fannie and John Hertz Foundation. B.T. acknowledges funding from the National Science Foundation Graduate Research Fellowship under Grant No. 2141064. L.G. acknowledges funding from the National Institutes of Health under Grant. No. T32GM087237. We are grateful to gifts from Open Philanthropy Project and the Aphorism Foundation (to K.M.E.). We thank Pranam Chatterjee and Sam Goldman for feedback on a draft of this manuscript. All figures were made with BioRender.

## Author contributions

V.S., B.T., and K.M.E. conceived the study. V.S. designed and implemented FLIGHTED and B.T. developed DHARMA and gathered the experimental data, both under the supervision of K.M.E.L.G. aided in benchmarking and optimization of FLIGHTED-DHARMA. V.S. wrote the first draft of the paper. All authors contributed to editing and approved the submission.

## References

1. Johnston, K. E., Fannjiang, C., Wittmann, B. J., Hie, B. L., Yang, K. K., and Wu, Z. Machine Learning for Protein Engineering (2023). http://arxiv.org/abs/2305.16634. doi:10.48550/arXiv.2305.16634 arXiv:2305.16634 [q-bio].

2. Frappier, V., and Keating, A. E. (2021). Data-driven computational protein design. Current Opinion in Structural Biology 69, 63–69. https://doi.org/10.1016/j.sbi.2021.03.009. doi:10.1016/J.SBI.2021.03.009. Publisher: Elsevier Ltd.

3. Bryant, D. H., Bashir, A., Sinai, S., Jain, N. K., Ogden, P. J., Riley, P. F., Church, G. M., Colwell, L. J., and Kelsic, E. D. (2021). Deep diversification of an AAV capsid protein by machine learning. Nature Biotechnology 39, 691–696. http://www.nature.com/articles/s41587-020-00793-4. doi:10.1038/s41587-020-00793-4.

4. Biswas, S., Khimulya, G., Alley, E. C., Esvelt, K. M., and Church, G. M. (2021). Low-N protein engineering with data-efficient deep learning. Nature Methods 2021 18:4 18, 389–396. https://www.nature.com/articles/s41592-021-01100-y. doi:10.1038/s41592-021-01100-y. Publisher: Nature Publishing Group ISBN: 4159202101100.

5. Wu, Z., Jennifer Kan, S. B., Lewis, R. D., Wittmann, B. J., and Arnold, F. H. (2019). Machine learning-assisted directed protein evolution with combinatorial libraries. Proceedings of the National Academy of Sciences of the United States of America 116, 8852–8858. doi:10.1073/PNAS.1901979116/SUPPL_FILE/PNAS.1901979116.SAPP.PDF. ArXiv: 1902.07231 Publisher: National Academy of Sciences.

6. Li, F.-Z., Amini, A. P., Yue, Y., Yang, K. K., and Lu, A. X. Feature Reuse and Scaling: Understanding Transfer Learning with Protein Language Models (2024). https://www.biorxiv.org/content/10.1101/2024.02.05.578959v1.full.pdf.

7. Dallago, C., Mou, J., Johnston, K. E., Wittmann, B. J., Bhattacharya, N., Goldman, S., Madani, A., and Yang, K. K. FLIP: Benchmark tasks in fitness landscape inference for proteins (2022). https://www.biorxiv.org/content/10.1101/2021.11.09.467890v2. doi:10.1101/2021.11.09.467890 section: New Results Type: article.

8. Rives, A., Goyal, S., Meier, J., Guo, D., Ott, M., Zitnick, C. L., Ma, J., and Fergus, R. (2021). Biological structure and function emerge from scaling unsupervised learning to 250 million protein sequences. Proceedings of the National Academy of Sciences 118, 622803. https://doi.org/10.1101/622803. doi:10.1101/622803. Publisher: bioRxiv.

9. Rao, R., Liu, J., Verkuil, R., Meier, J., Canny, J. F., Abbeel, P., Sercu, T., and Rives, A. MSA Transformer (2021). https://www.biorxiv.org/content/10.1101/2021.02.12.430858v3. doi:10.1101/2021.02.12.430858 section: New Results Type: article.

10. Nijkamp, E., Ruffolo, J., Weinstein, E. N., Naik, N., and Madani, A. ProGen2: Exploring the Boundaries of Protein Language Models (2022). http://arxiv.org/abs/2206.13517 arXiv:2206.13517 [cs, q-bio].

11. Notin, P., Dias, M., Frazer, J., Marchena-Hurtado, J., Gomez, A., Marks, D. S., and Gal, Y. Tranception: protein fitness prediction with autoregressive transformers and inference-time retrieval. In: Proceedings of the 39th International Conference on Machine Learning. arXiv (2022):http://arxiv.org/abs/2205.13760 number: arXiv:2205.13760 arXiv:2205.13760 [cs].

12. Fernandez-de Cossio-Diaz, J., Uguzzoni, G., and Pagnani, A. (2021). Unsupervised Inference of Protein Fitness Landscape from Deep Mutational Scan. Molecular Biology and Evolution 38, 318–328. https://academic.oup.com/mbe/article/38/1/318/5889958. doi:10.1093/molbev/msaa204. Publisher: Oxford University Press.

13. Rubin, A. F., Gelman, H., Lucas, N., Bajjalieh, S. M., Papenfuss, A. T., Speed, T. P., and Fowler, D. M. (2017). A statistical framework for analyzing deep mutational scanning data. Genome Biology 18, 150. https://doi.org/10.1186/s13059-017-1272-5. doi:10.1186/s13059-017-1272-5.

14. Busia, A., and Listgarten, J. Model-based differential sequencing analysis (2023). http://biorxiv.org/lookup/doi/10.1101/2023.03.29.534803. doi:10.1101/2023.03.29.534803.

15. Sesta, L., Pagnani, A., Fernandez-de Cossio-Diaz, J., and Uguzzoni, G. (2024). Inference of annealed protein fitness landscapes with AnnealDCA. PLOS Computational Biology 20, e1011812. https://journals.plos.org/ploscompbiol/article?id=10.1371/journal.pcbi.1011812. doi:10.1371/journal.pcbi.1011812. Publisher: Public Library of Science.

16. Zhu, D., Brookes, D. H., Busia, A., Carneiro, A., Fannjiang, C., Popova, G., Shin, D., Donohue, K. C., Lin, L. F., Miller, Z. M., Williams, E. R., Chang, E. F., Nowakowski, T. J., Listgarten, J., and Schaffer, D. V. (2024). Optimal trade-off control in machine learning–based library design, with application to adeno-associated virus (AAV) for gene therapy. Science Advances 10, eadj3786. https://www.science.org/doi/10.1126/sciadv.adj3786. doi:10.1126/sciadv.adj3786. Publisher: American Association for the Advancement of Science.

17. Tu, B., Sundar, V., and Esvelt, K. An ultra-high-throughput method for measuring biomolecular activities (2024). https://www.biorxiv.org/content/10.1101/2022.03.09.483646v4. doi:10.1101/2022.03.09.483646 pages: 2022.03.09.483646 Section: New Results.

18. Hoffman, M., Blei, D. M., Wang, C., and Paisley, J. Stochastic Variational Inference (2013). http://arxiv.org/abs/1206.7051. doi:10.48550/arXiv.1206.7051 arXiv:1206.7051 [cs, stat].

19. Kingma, D. P., and Welling, M. Auto-Encoding Variational Bayes (2014). http://arxiv.org/abs/1312.6114 number: arXiv:1312.6114 arXiv:1312.6114 [cs, stat].

20. Sohn, K., Lee, H., and Yan, X. Learning Structured Output Representation using Deep Conditional Generative Models. In: Advances in Neural Information Processing Systems vol. 28. Curran Associates, Inc. (2015):https://papers.nips.cc/paper_files/paper/2015/hash/8d55a249e6baa5c06772297520da2051-Abstract.html.

21. Mandecki, W., Chen, J. Y.-C., and Grihalde, N. (1995). A Mathematical Model for Biopanning (Affinity Selection) Using Peptide Libraries on Filamentous Phage. Journal of Theoretical Biology 176, 523–530. https://linkinghub.elsevier.com/retrieve/pii/S0022519385702184. doi:10.1006/jtbi.1995.0218.

22. Russel, M., Lowman, H. B., and Clackson, T. Introduction to phage biology and phage display. In: Phage Display: A Practical Approach (26). Practical Approach Series Oxford: Oxford University Press. ISBN 978-0-19-963873-4 (2004):(26).

23. Matuszewski, S., Hildebrandt, M. E., Ghenu, A.-H., Jensen, J. D., and Bank, C. (2016). A Statistical Guide to the Design of Deep Mutational Scanning Experiments. Genetics 204, 77–87. https://doi.org/10.1534/genetics.116.190462. doi:10.1534/genetics.116.190462.

24. Wu, N. C., Dai, L., Olson, C. A., Lloyd-Smith, J. O., and Sun, R. (2016). Adaptation in protein fitness landscapes is facilitated by indirect paths. eLife 5. doi:10.7554/eLife.16965. Publisher: eLife Sciences Publications Ltd.

25. Miller, S. M., Wang, T., and Liu, D. R. (2020). Phage-assisted continuous and non-continuous evolution. Nature Protocols 2020 15:12 15, 4101–4127. https://www.nature.com/articles/s41596-020-00410-3. doi:10.1038/s41596-020-00410-3. Publisher: Nature Publishing Group.

26. Raj, A., and Oudenaarden, A. v. (2008). Nature, Nurture, or Chance: Stochastic Gene Expression and Its Consequences. Cell 135, 216–226. https://www.cell.com/cell/abstract/S0092-8674(08)01243-9. doi:10.1016/j.cell.2008.09.050. Publisher: Elsevier.

27. Meier, J., Rao, R., Verkuil, R., Liu, J., Sercu, T., and Rives, A. (2021). Language models enable zero-shot prediction of the effects of mutations on protein function. Proceedings of the 35th Conference on Neural Information Processing Systems.

28. Yang, K. K., Lu, A. X., and Fusi, N. Convolutions are competitive with transformers for protein sequence pretraining (2022). https://www.biorxiv.org/content/10.1101/2022.05.19.492714v1.full.pdf.

29. Hsu, C., Nisonoff, H., Fannjiang, C., and Listgarten, J. (2022). Learning protein fitness models from evolutionary and assay-labeled data. Nature Biotechnology. https://www.nature.com/articles/s41587-021-01146-5.

30. Notin, P., Kollasch, A. W., Ritter, D., Van Niekerk, L., Paul, S., Spinner, H., Rollins, N., Shaw, A., Weitzman, R., Frazer, J., Dias, M., Franceschi, D., Orenbuch, R., Gal, Y., and Marks, D. S. ProteinGym: Large-Scale Benchmarks for Protein Design and Fitness Prediction. In: NeurIPS 2023 Track on Datasets and Benchmarks (2023):http://biorxiv.org/lookup/doi/10.1101/2023.12.07.570727. doi:10.1101/2023.12.07.570727.

31. Curran, P. A. Monte Carlo error analyses of Spearman’s rank test (2015). http://arxiv.org/abs/1411.3816 arXiv:1411.3816 [astro-ph, physics:physics, stat].

32. Lin, Z., Akin, H., Rao, R., Hie, B., Zhu, Z., Lu, W., Fazel-Zarandi, M., Sercu, T., Candido, S., and Rives, A. Language models of protein sequences at the scale of evolution enable accurate structure prediction (2022). https://www.biorxiv.org/content/10.1101/2022.07.20.500902v1.full.pdf.

33. Bingham, E., Chen, J. P., Jankowiak, M., Obermeyer, F., Pradhan, N., Karaletsos, T., Singh, R., Szerlip, P., Horsfall, P., and Goodman, N. D. (2019). Pyro: Deep Universal Probabilistic Programming. Journal of Machine Learning Research 20, 1–6.

34. Paszke, A., Gross, S., Massa, F., Lerer, A., Bradbury, J., Chanan, G., Killeen, T., Lin, Z., Gimelshein, N., Antiga, L., Desmaison, A., Kopf, A., Yang, E., DeVito, Z., Raison, M., Tejani, A., Chilamkurthy, S., Steiner, B., Fang, L., Bai, J., and Chintala, S. PyTorch: An Imperative Style, High-Performance Deep Learning Library. In: 33rd Conference on Neural Information Processing Systems (2019):.

35. Jones, E., Oliphant, T., Peterson, P., and Others. SciPy: Open source scientific tools for Python (2001). http://www.scipy.org/.

36. Rao, R., Bhattacharya, N., Thomas, N., Duan, Y., Chen, X., Canny, J., Abbeel, P., and Song, Y. S. Evaluating protein transfer learning with TAPE. In: Advances in Neural Information Processing Systems (2019):https://www.biorxiv.org/content/10.1101/676825v1. doi:10.1101/676825 arXiv: 1906.08230 ISSN: 1049-5258.

37. Elnaggar, A., Heinzinger, M., Dallago, C., Rehawi, G., Wang, Y., Jones, L., Gibbs, T., Feher, T., Angerer, C., Steinegger, M., Bhowmik, D., and Rost, B. (2021). ProtTrans: Towards Cracking the Language of Life’s Code Through Self-Supervised Learning. IEEE Transactions on Pattern Analysis and Machine Intelligence 14. http://biorxiv.org/lookup/doi/10.1101/2020.07.12.199554. doi:10.1101/2020.07.12.199554.

38. Hopf, T. A., Ingraham, J. B., Poelwijk, F. J., Schärfe, C. P. I., Springer, M., Sander, C., and Marks, D. S. (2017). Mutation effects predicted from sequence co-variation. Nature Biotechnology 35, 128–135. doi:10.1038/nbt.3769.

