## Supplemental Information for "FLIGHTED: Inferring Fitness Landscapes from Noisy High-Throughput Experimental Data"

### Impact of Read Count on Fitnesses in the TEV Landscape

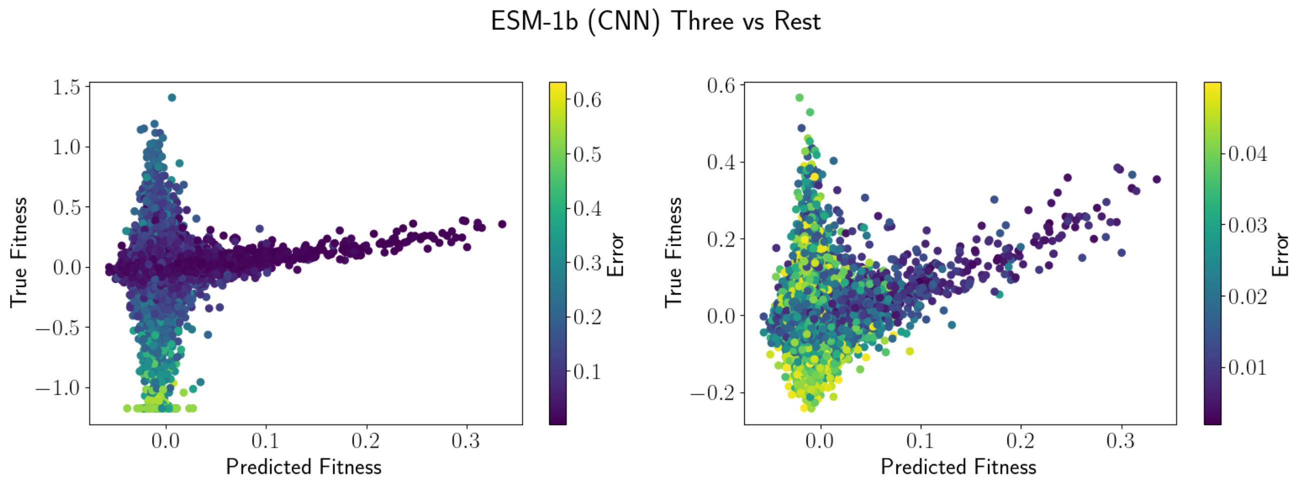


Figure S1: Sample Model Predictions on TEV Dataset, produced by ESM-1b CNN on three-vs-rest. Left includes all data points and right includes data points filtered with error $< 0.05$ to highlight most accurate model predictions.

We begin by examining model predictions for a single high-performing model (the ESM-1b CNN on three-vs-rest) in Figure S1; here, predicted fitness refers to the fitness predicted by the ESM-1b model, and true fitness refers to the FLIGHTED-measured fitness with the error (variance) shown on the colorbar. The model has reasonably high accuracy on predictions with very low variance ($< 0.01$) and predicts most other data points with high variance to be inactive, suggesting the model has learned to filter by variance when making predictions. Supplementary Figure S6d suggests that the primary determinant of variance is read count, so this implies the model has learned read count on the test set, potentially surprising.

We investigated this result further to ensure that there was no accidental data leakage at any point. There are two effects we identified that allow the model to learn read count on the test set. First, the T7 lysozyme is slightly toxic to the cell and when not cleaved (i.e. when the protease is inactive), results in a lower read count. In Figure S2, we compare the read count in libraries with and without the lysozyme and see that active variants have higher read count in the library with the lysozyme. Since the models are trained to learn fitness, they can also implicitly learn about this effect.

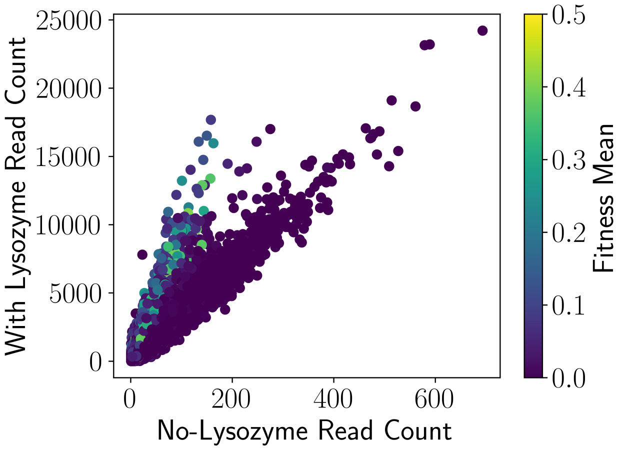


Figure S2: Read Count With and Without T7 Lysozyme. Active variants have higher read counts with the T7 lysozyme.

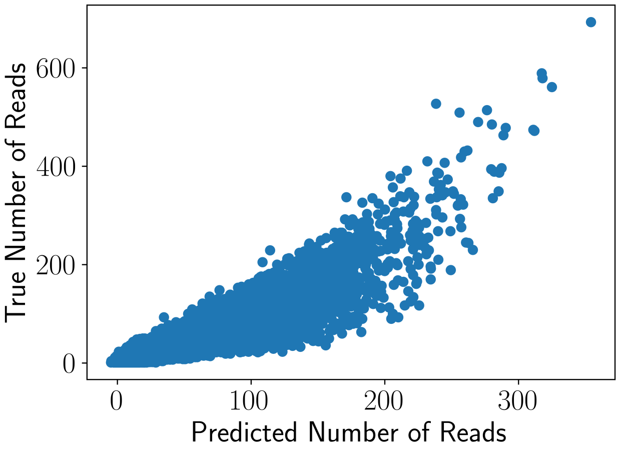


Figure S3: Control Model Trained on No-Lysozyme Read Count, showing that read count is learnable.

Second, the initial (no-lysozyme) library construction was generated by a potentially biased synthesis process, so the initial read count is learnable, i.e. similar variants have similar read counts. We tested this by training a model (the same ESM-1b CNN) on the initial read counts of a training set consisting of triple mutants and predicting read counts on the quadruple mutants, as shown in Figure S3. Since read count is learnable, we conclude that models are learning a combination of the initial read count from the biased library and TEV protease fitness when making predictions. This dataset and these models are still generally useful for predicting TEV protease activity and benchmarking ML models, but any active variant that had low read count in the initial library would likely be a false negative on any model trained on this dataset. To confirm that the models did learn TEV protease activity, we verified that the models predicted low activity for variants with high read count and low activity, as expected. In the future, we aim to reduce library bias to generate higher-quality TEV protease datasets and more accurate models to mitigate this issue.

| Model | One vs Rest | One vs Rest FLIGHTED | Two vs Rest | Two vs Rest FLIGHTED |
|---|---|---|---|---|
| Linear | 1.4706 ± 0.0127 | 12.0096 ± 0.0709 | 1.5174 ± 0.0020 | 9.8847 ± 0.0342 |
| TAPE | 1.5062 ± 0.0090 | 11.8502 ± 0.2601 | 1.9685 ± 0.1161 | 7.1776 ± 0.3339 |
| ESM-1b | 1.4898 ± 0.0004 | **11.2226 ± 0.0750** | 1.4427 ± 0.0107 | 5.7642 ± 0.2590 |
| ESM-1v | 1.5132 ± 0.0016 | 14.7296 ± 0.0125 | 1.5646 ± 0.0153 | 6.7795 ± 0.2354 |
| ESM-2 (8M) | 1.5169 ± 0.0177 | 11.6852 ± 0.0226 | 1.4388 ± 0.0081 | 7.6910 ± 0.1750 |
| ESM-2 (35M) | 1.5218 ± 0.0218 | 11.5808 ± 0.0899 | 1.4034 ± 0.0103 | 8.2036 ± 0.0373 |
| ESM-2 (150M) | 1.5129 ± 0.0210 | 11.5738 ± 0.1378 | 1.4099 ± 0.0057 | 8.5423 ± 0.0238 |
| ESM-2 (650M) | 1.4963 ± 0.0024 | 11.5768 ± 0.2312 | 1.4285 ± 0.0288 | 5.5143 ± 0.2098 |
| ESM-2 | 1.9206 ± 0.0334 | 12.0280 ± 0.1538 | 1.3472 ± 0.0071 | 5.0572 ± 0.3065 |
| ProtT5 | 1.5288 ± 0.0148 | 11.6078 ± 0.0677 | 1.3567 ± 0.0259 | 5.4940 ± 0.0949 |
| CARP (600k) | 1.5033 ± 0.0132 | 11.6508 ± 0.0983 | 1.5659 ± 0.0472 | 11.2536 ± 0.1272 |
| CARP (38M) | 1.5139 ± 0.0088 | 11.6778 ± 0.1284 | 1.5768 ± 0.0059 | 11.0896 ± 0.1535 |
| CARP (76M) | 1.4884 ± 0.0145 | 11.5002 ± 0.2140 | 1.5607 ± 0.0031 | 9.1447 ± 0.1564 |
| CARP | 1.5070 ± 0.0082 | 11.7667 ± 0.2218 | 1.5402 ± 0.0041 | 9.4377 ± 0.2619 |
| CNN | 2.0506 ± 0.2307 | 13.8411 ± 1.6964 | 1.3523 ± 0.0539 | 4.0959 ± 0.1426 |
| TAPE (CNN) | 2.0937 ± 0.1898 | 15.5570 ± 0.5312 | 1.1251 ± 0.0631 | 4.9553 ± 0.6760 |
| ESM-1b (CNN) | 1.8592 ± 0.1246 | 14.7160 ± 1.1399 | 1.1482 ± 0.0239 | 4.6976 ± 0.4697 |
| ESM-1v (CNN) | 2.4676 ± 0.2503 | 15.0658 ± 1.1423 | 1.1188 ± 0.0212 | 5.6638 ± 0.6273 |
| ESM-2 (8M, CNN) | 2.3919 ± 0.1627 | 15.5196 ± 0.4909 | 1.0913 ± 0.0416 | 4.8430 ± 0.5947 |
| ESM-2 (35M, CNN) | 2.4945 ± 0.4818 | 14.7227 ± 0.6925 | 1.0381 ± 0.0769 | 3.8103 ± 0.1281 |
| ESM-2 (150M, CNN) | 2.2499 ± 0.5033 | 15.6027 ± 1.0554 | 1.1085 ± 0.1243 | 4.7428 ± 0.4609 |
| ESM-2 (650M, CNN) | 1.8934 ± 0.3340 | 14.2969 ± 0.7712 | 1.1476 ± 0.0261 | 4.7025 ± 0.6881 |
| ESM-2 (CNN) | 1.9511 ± 0.1986 | 14.2011 ± 1.6117 | 1.3850 ± 0.0382 | 5.3430 ± 0.8577 |
| ProtT5 (CNN) | 1.8554 ± 0.0745 | 14.9394 ± 0.4143 | 1.0373 ± 0.0246 | 5.0151 ± 1.0703 |
| CARP (600k, CNN) | 2.6833 ± 0.3544 | 14.1731 ± 0.3073 | 1.2257 ± 0.0350 | 5.2804 ± 0.9685 |
| CARP (38M, CNN) | 2.0301 ± 0.2404 | 15.0381 ± 1.5099 | 1.1312 ± 0.0228 | 4.6168 ± 0.4402 |
| CARP (76M, CNN) | 1.9932 ± 0.0644 | 14.6097 ± 0.3420 | 1.1944 ± 0.0091 | 3.8172 ± 0.2085 |
| CARP (CNN) | 1.7704 ± 0.0174 | 15.0078 ± 0.8024 | 1.3109 ± 0.0596 | 5.4695 ± 0.7086 |
| ESM-1v (Augmented) | 1.4845 ± 0.0018 | 11.7308 ± 0.2460 | 1.5685 ± 0.0150 | 10.0828 ± 0.1070 |
| ESM-2 (8M, Augmented) | 1.5088 ± 0.0170 | 11.9113 ± 0.0402 | 1.6641 ± 0.0156 | 10.0938 ± 0.0589 |
| ESM-2 (35M, Augmented) | 1.5102 ± 0.0220 | 11.9370 ± 0.0565 | 1.6291 ± 0.0127 | 10.1836 ± 0.1647 |
| ESM-2 (150M, Augmented) | 1.5230 ± 0.0233 | 11.9377 ± 0.1009 | 1.5985 ± 0.0040 | 9.7887 ± 0.1380 |
| ESM-2 (650M, Augmented) | 1.4985 ± 0.0086 | 11.8261 ± 0.0717 | 1.6410 ± 0.0120 | 9.6624 ± 0.1501 |
| ESM-2 (Augmented) | 1.5115 ± 0.0318 | 11.7135 ± 0.1641 | 1.6180 ± 0.0112 | 9.6967 ± 0.0494 |
| CARP (600k, Augmented) | **1.4652 ± 0.0021** | 11.9570 ± 0.0983 | 1.5064 ± 0.0039 | 9.9571 ± 0.1009 |
| CARP (38M, Augmented) | 1.4777 ± 0.0186 | 11.9582 ± 0.0450 | 1.4982 ± 0.0025 | 9.9154 ± 0.0556 |
| CARP (76M, Augmented) | 1.4757 ± 0.0199 | 11.7828 ± 0.1579 | 1.4985 ± 0.0058 | 9.8734 ± 0.0189 |
| CARP (Augmented) | 1.4660 ± 0.0022 | 12.0082 ± 0.0337 | 1.5083 ± 0.0061 | 9.9470 ± 0.0716 |
| EVMutation (Augmented) | 1.4708 ± 0.0093 | 11.6674 ± 0.3938 | 1.4526 ± 0.0022 | 7.0896 ± 0.0080 |
| ESM-2 (8M, Finetuned) | 1.5318 ± 0.0264 | 14.9039 ± 2.5033 | 1.6052 ± 0.0357 | 4.7209 ± 0.2025 |
| ESM-2 (35M, Finetuned) | 1.5500 ± 0.0967 | 13.5396 ± 0.3245 | 1.3019 ± 0.0551 | 3.7613 ± 0.0887 |
| ESM-2 (150M, Finetuned) | 1.5245 ± 0.0026 | 11.4452 ± 0.2846 | 1.1094 ± 0.0467 | **3.3694 ± 0.1247** |
| ESM-2 (8M, CNN, Finetuned) | 2.0855 ± 0.3826 | 15.5261 ± 0.3309 | 1.0087 ± 0.0216 | 4.5865 ± 0.4793 |
| ESM-2 (35M, CNN, Finetuned) | 2.2368 ± 0.4315 | 13.8723 ± 0.8406 | **0.8978 ± 0.0305** | 3.5521 ± 0.3629 |
| ESM-2 (150M, CNN, Finetuned) | 2.8614 ± 0.2362 | 14.9521 ± 0.7068 | 1.0092 ± 0.1637 | 3.9133 ± 0.2355 |

Table S1: Raw Model Performance on GB1 Datasets (one-vs-rest and two-vs-rest only). See Methods for a detailed description of all models and dataset splits. Bolded models are the highest-performing in a given task. Highlighted models are not statistically significantly different via a two-sided t-test from the top model at a significance level of $p < 0.05$.

| Model | Three vs Rest | Three vs Rest FLIGHTED | Low vs High | Low vs High FLIGHTED |
|---|---|---|---|---|
| Linear | 1.5172 ± 0.0049 | 8.1622 ± 0.0122 | 4.6324 ± 0.0126 | 7.5193 ± 0.1734 |
| TAPE | 0.9591 ± 0.0295 | 3.6993 ± 0.0375 | 3.5252 ± 0.0394 | 2.4557 ± 0.1641 |
| ESM-1b | 0.9484 ± 0.0104 | 3.2003 ± 0.0183 | 3.5314 ± 0.0218 | 2.5016 ± 0.1014 |
| ESM-1v | 0.9261 ± 0.0185 | 3.4494 ± 0.0882 | 3.4070 ± 0.0201 | 1.7771 ± 0.0768 |
| ESM-2 (8M) | 1.0639 ± 0.0142 | 4.3834 ± 0.0081 | 3.6068 ± 0.0302 | 3.4855 ± 0.0535 |
| ESM-2 (35M) | 0.9534 ± 0.0022 | 3.8483 ± 0.0362 | 3.5095 ± 0.0738 | 2.9552 ± 0.0789 |
| ESM-2 (150M) | 0.9726 ± 0.0052 | 4.3019 ± 0.1824 | 3.5763 ± 0.0271 | 3.0476 ± 0.0451 |
| ESM-2 (650M) | 0.7834 ± 0.0036 | 2.9651 ± 0.0779 | 3.4553 ± 0.0089 | 2.3014 ± 0.0535 |
| ESM-2 | 0.8850 ± 0.0038 | 2.8603 ± 0.0090 | 3.4237 ± 0.0502 | 2.1913 ± 0.0138 |
| ProtT5 | 0.8706 ± 0.0072 | 3.0169 ± 0.0259 | 3.3162 ± 0.0289 | 2.0257 ± 0.0282 |
| CARP (600k) | 1.1682 ± 0.0112 | 5.5622 ± 0.0866 | 3.9904 ± 0.0258 | 4.4635 ± 0.1299 |
| CARP (38M) | 1.3215 ± 0.0152 | 6.4299 ± 0.1360 | 4.2556 ± 0.0272 | 5.4571 ± 0.0665 |
| CARP (76M) | 1.1813 ± 0.0074 | 5.3000 ± 0.1357 | 4.1310 ± 0.0059 | 4.7139 ± 0.2040 |
| CARP | 1.2078 ± 0.0052 | 5.0631 ± 0.2182 | 4.1793 ± 0.0399 | 5.0080 ± 0.1777 |
| CNN | 0.5158 ± 0.0101 | 1.5133 ± 0.0458 | 3.1096 ± 0.0319 | **0.6077 ± 0.0747** |
| TAPE (CNN) | 0.4504 ± 0.0039 | 1.2584 ± 0.0790 | 3.1693 ± 0.0720 | 0.7386 ± 0.0647 |
| ESM-1b (CNN) | 0.4406 ± 0.0066 | 1.2828 ± 0.0583 | 3.2526 ± 0.1097 | 0.8580 ± 0.1646 |
| ESM-1v (CNN) | **0.4218 ± 0.0148** | 1.0939 ± 0.0647 | 3.2323 ± 0.1479 | 0.8102 ± 0.0425 |
| ESM-2 (8M, CNN) | 0.4587 ± 0.0071 | 1.3937 ± 0.1154 | **3.1004 ± 0.0596** | 0.8299 ± 0.0669 |
| ESM-2 (35M, CNN) | 0.4736 ± 0.0061 | 1.3928 ± 0.0585 | 3.1440 ± 0.0846 | 0.6806 ± 0.1123 |
| ESM-2 (150M, CNN) | 0.4724 ± 0.0051 | 1.3727 ± 0.0349 | 3.1400 ± 0.0445 | 0.7269 ± 0.1354 |
| ESM-2 (650M, CNN) | 0.4396 ± 0.0515 | 1.2475 ± 0.1657 | 3.3286 ± 0.0471 | 0.8302 ± 0.0805 |
| ESM-2 (CNN) | 0.4440 ± 0.0259 | 1.2242 ± 0.1160 | 3.3670 ± 0.1110 | 0.8903 ± 0.0651 |
| ProtT5 (CNN) | 0.4576 ± 0.0232 | 1.1711 ± 0.0338 | 3.2488 ± 0.0403 | 0.8296 ± 0.0278 |
| CARP (600k, CNN) | 0.5965 ± 0.0216 | 1.7806 ± 0.0977 | 3.2274 ± 0.0671 | 0.9752 ± 0.1267 |
| CARP (38M, CNN) | 0.4854 ± 0.0094 | 1.1501 ± 0.0552 | 3.1777 ± 0.0413 | 0.8558 ± 0.0859 |
| CARP (76M, CNN) | 0.5070 ± 0.0409 | 1.2360 ± 0.0822 | 3.2070 ± 0.0067 | 0.8577 ± 0.0944 |
| CARP (CNN) | 0.4661 ± 0.0215 | 1.3619 ± 0.1444 | 3.2760 ± 0.0515 | 0.7709 ± 0.1090 |
| ESM-1v (Augmented) | 1.4944 ± 0.0063 | 7.9110 ± 0.1260 | 4.6129 ± 0.0221 | 7.1360 ± 0.1249 |
| ESM-2 (8M, Augmented) | 1.5203 ± 0.0064 | 7.9353 ± 0.0718 | 4.6053 ± 0.0032 | 7.3604 ± 0.1191 |
| ESM-2 (35M, Augmented) | 1.4982 ± 0.0024 | 7.9133 ± 0.0427 | 4.6106 ± 0.0130 | 7.3177 ± 0.0638 |
| ESM-2 (150M, Augmented) | 1.4950 ± 0.0080 | 7.6113 ± 0.0829 | 4.5462 ± 0.0280 | 6.9321 ± 0.0473 |
| ESM-2 (650M, Augmented) | 1.5175 ± 0.0049 | 7.3998 ± 0.1201 | 4.5873 ± 0.0184 | 6.8160 ± 0.0256 |
| ESM-2 (Augmented) | 1.5265 ± 0.0051 | 7.7374 ± 0.1413 | 4.6023 ± 0.0446 | 7.1383 ± 0.0245 |
| CARP (600k, Augmented) | 1.5168 ± 0.0064 | 8.0217 ± 0.0664 | 4.5907 ± 0.0208 | 7.4045 ± 0.1267 |
| CARP (38M, Augmented) | 1.5153 ± 0.0088 | 8.0537 ± 0.0542 | 4.5971 ± 0.0493 | 7.4798 ± 0.0527 |
| CARP (76M, Augmented) | 1.5177 ± 0.0027 | 8.2109 ± 0.1684 | 4.6521 ± 0.0142 | 7.5263 ± 0.0519 |
| CARP (Augmented) | 1.5186 ± 0.0034 | 8.2201 ± 0.0617 | 4.6398 ± 0.0729 | 7.5125 ± 0.1570 |
| EVMutation (Augmented) | 1.5126 ± 0.0068 | 6.6964 ± 0.0356 | 4.5886 ± 0.0487 | 7.2021 ± 0.0813 |
| ESM-2 (8M, Finetuned) | 0.5035 ± 0.0127 | **0.9590 ± 0.0316** | 3.4907 ± 0.0767 | 1.0527 ± 0.0363 |
| ESM-2 (35M, Finetuned) | 0.5780 ± 0.0566 | 1.1488 ± 0.0564 | 3.5533 ± 0.0036 | 1.0101 ± 0.0532 |
| ESM-2 (150M, Finetuned) | 0.4928 ± 0.0451 | 1.0775 ± 0.1876 | 3.6195 ± 0.0081 | 1.0493 ± 0.0379 |
| ESM-2 (8M, CNN, Finetuned) | 0.4613 ± 0.0162 | 1.0441 ± 0.0517 | 3.2580 ± 0.0426 | 0.9389 ± 0.0713 |
| ESM-2 (35M, CNN, Finetuned) | 0.4359 ± 0.0166 | 1.0510 ± 0.0885 | 3.2956 ± 0.0378 | 0.7850 ± 0.1027 |
| ESM-2 (150M, CNN, Finetuned) | 0.4476 ± 0.0211 | 1.0069 ± 0.0271 | 3.2438 ± 0.0467 | 0.6934 ± 0.0123 |

Table S2: Raw Model Performance on GB1 Datasets (three-vs-rest and low-vs-high only). See Methods for a detailed description of all models and dataset splits. Bolded models are the highest-performing in a given task. Highlighted models are not statistically significantly different via a two-sided t-test from the top model at a significance level of $p < 0.05$.

| Model | One vs Rest | Two vs Rest | Three vs Rest |
|---|---|---|---|
| Linear | 526.0933 ± 22.7232 | 473.0123 ± 30.1776 | 61.9535 ± 9.9049 |
| TAPE | 297.5274 ± 16.9196 | 42.1769 ± 5.7037 | 10.1017 ± 2.9131 |
| ESM-1b | 165.9262 ± 0.8583 | 19.2814 ± 2.1384 | 4.7189 ± 0.0789 |
| ESM-1v | **96.5753 ± 4.8990** | 31.6219 ± 8.8151 | 5.0249 ± 0.2268 |
| ESM-2 (8M) | 487.1830 ± 67.0827 | 77.3104 ± 13.6258 | 26.0598 ± 11.2005 |
| ESM-2 (35M) | 201.0534 ± 14.7650 | 38.3270 ± 14.7239 | 5.3898 ± 0.0861 |
| ESM-2 (150M) | 253.0745 ± 17.7263 | 21.2191 ± 4.1567 | 5.8834 ± 0.7763 |
| ESM-2 (650M) | 129.0170 ± 3.0636 | 15.7421 ± 2.0325 | 5.0022 ± 0.5990 |
| ESM-2 | 455.1378 ± 8.2257 | 21.5850 ± 7.1289 | 16.4862 ± 17.6678 |
| ProtT5 | 128.9400 ± 5.0140 | 16.2017 ± 2.1542 | 4.5883 ± 0.2502 |
| CARP (600k) | 370.3106 ± 61.6088 | 67.1536 ± 5.6832 | 17.8028 ± 0.6229 |
| CARP (38M) | 779.2183 ± 6.6225 | 69.2424 ± 6.2080 | 11.5826 ± 1.0835 |
| CARP (76M) | 293.7587 ± 52.0933 | 59.6290 ± 2.5587 | 14.5622 ± 0.9285 |
| CARP | 661.4456 ± 227.2850 | 201.7350 ± 12.4057 | 25.1076 ± 8.3689 |
| CNN | 613.7185 ± 242.4505 | 169.1024 ± 7.8148 | 6.4333 ± 0.1398 |
| TAPE (CNN) | 535.2333 ± 132.6787 | 119.1340 ± 10.3364 | 5.5736 ± 0.2825 |
| ESM-1b (CNN) | 565.8093 ± 203.6468 | 40.1009 ± 9.9084 | 4.5211 ± 0.0431 |
| ESM-1v (CNN) | 678.3326 ± 164.8031 | 28.9555 ± 0.4254 | 4.7168 ± 0.1845 |
| ESM-2 (8M, CNN) | 537.5033 ± 238.1350 | 56.3152 ± 9.9305 | 5.3183 ± 0.1928 |
| ESM-2 (35M, CNN) | 299.9104 ± 14.8647 | 26.8592 ± 5.2674 | 224.8132 ± 191.9686 |
| ESM-2 (150M, CNN) | 676.5742 ± 103.0750 | 31.0988 ± 5.3971 | 4.7040 ± 0.1121 |
| ESM-2 (650M, CNN) | 915.3362 ± 23.6130 | 66.0526 ± 3.5709 | 4.8076 ± 0.0419 |
| ESM-2 (CNN) | 1361.5717 ± 539.0819 | 87.6785 ± 10.3623 | 4.9448 ± 0.1656 |
| ProtT5 (CNN) | 900.8455 ± 233.7611 | 61.6325 ± 3.8367 | 4.7374 ± 0.0591 |
| CARP (600k, CNN) | 479.3737 ± 9.2137 | 69.0239 ± 4.6387 | 6.8296 ± 0.4412 |
| CARP (38M, CNN) | 685.8954 ± 281.9056 | 79.6063 ± 9.2690 | 4.8927 ± 0.0288 |
| CARP (76M, CNN) | 581.1124 ± 86.5494 | 64.4380 ± 2.4957 | 4.7728 ± 0.1322 |
| CARP (CNN) | 797.2115 ± 459.2779 | 40.6028 ± 0.7333 | 4.6357 ± 0.1348 |
| ESM-1v (Augmented) | 238.4065 ± 5.2917 | 480.2571 ± 33.7659 | 63.6243 ± 3.8785 |
| ESM-2 (8M, Augmented) | 337.0354 ± 13.8129 | 492.2340 ± 28.7706 | 69.7858 ± 4.0155 |
| ESM-2 (35M, Augmented) | 216.1071 ± 23.8610 | 472.1725 ± 53.4941 | 71.2082 ± 6.5696 |
| ESM-2 (150M, Augmented) | 222.0371 ± 4.0988 | 490.2639 ± 30.5448 | 61.4350 ± 7.4693 |
| ESM-2 (650M, Augmented) | 338.1440 ± 190.0764 | 481.7004 ± 19.7602 | 69.2233 ± 11.1288 |
| ESM-2 (Augmented) | 226.2928 ± 33.0988 | 507.1823 ± 49.9358 | 69.0547 ± 3.8129 |
| CARP (600k, Augmented) | 555.9357 ± 65.4261 | 469.3298 ± 3.0498 | 65.6611 ± 1.8785 |
| CARP (38M, Augmented) | 608.4704 ± 69.0148 | 449.0583 ± 18.4007 | 61.7768 ± 5.5423 |
| CARP (76M, Augmented) | 562.6250 ± 15.5729 | 464.0757 ± 13.7924 | 64.1665 ± 7.0719 |
| CARP (Augmented) | 508.6772 ± 74.7882 | 459.4699 ± 36.2165 | 66.3554 ± 4.3394 |
| EVMutation (Augmented) | 247.1387 ± 20.9441 | 492.7133 ± 31.5477 | 62.2780 ± 6.3277 |
| ESM-2 (8M, Finetuned) | 260.1472 ± 66.3833 | 7.6464 ± 1.6408 | 4.2765 ± 0.0316 |
| ESM-2 (35M, Finetuned) | 273.4673 ± 15.2778 | **7.5348 ± 0.3203** | 4.2672 ± 0.0318 |
| ESM-2 (8M, CNN, Finetuned) | 517.6429 ± 199.1142 | 19.5290 ± 1.4134 | 4.7530 ± 0.1151 |
| ESM-2 (35M, CNN, Finetuned) | 211.5166 ± 46.2842 | 10.7844 ± 0.5173 | **4.2564 ± 0.0468** |

Table S3: Raw Model Performance on TEV Datasets. See Methods for a detailed description of all models and dataset splits. Bolded models are the highest-performing in a given task. Highlighted models are not statistically significantly different via a two-sided t-test from the top model at a significance level of $p < 0.05$.

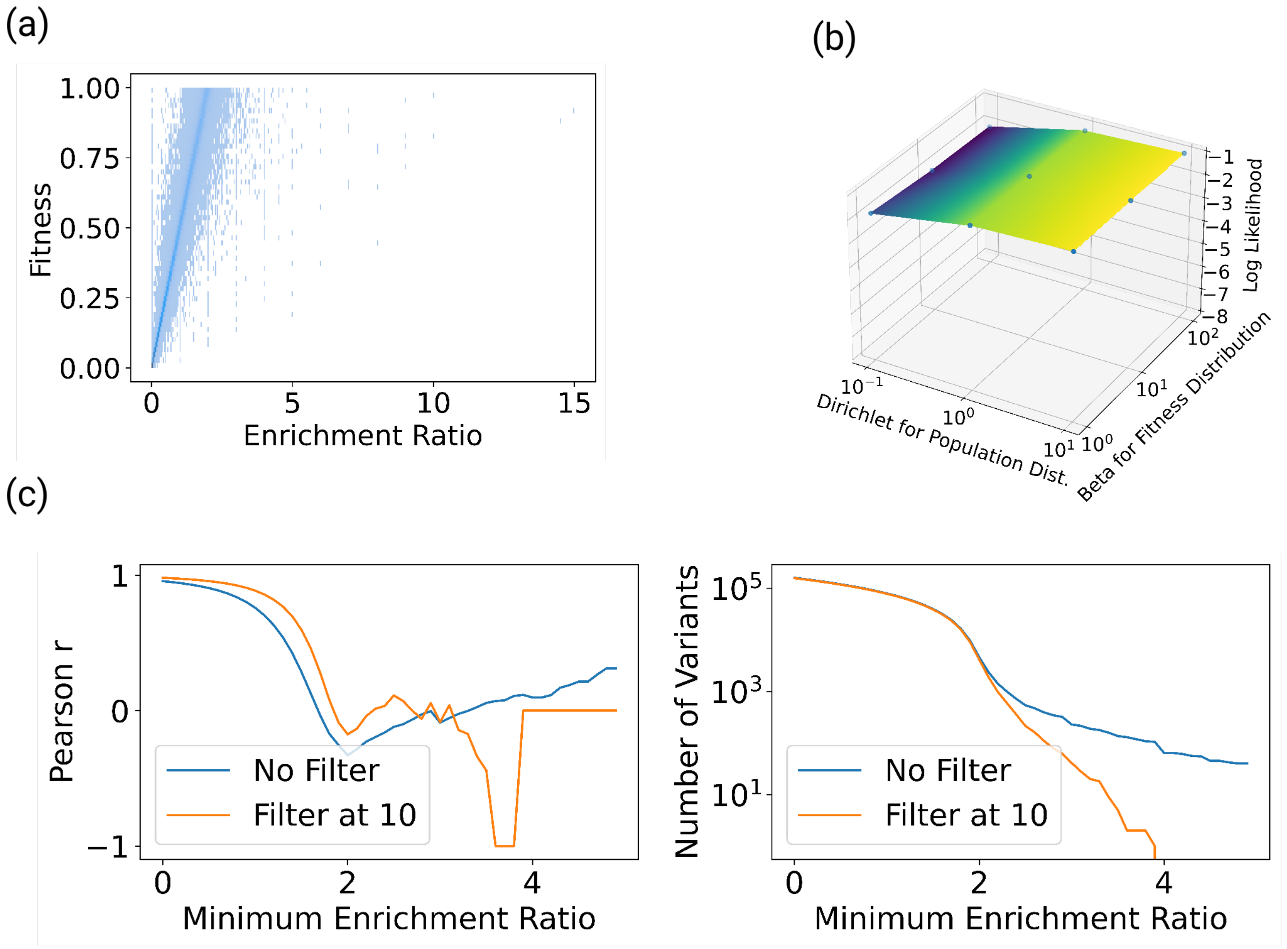


Figure S4: Noise in Single-Step Selection Simulations and Robustness of FLIGHTED-Selection Model Performance. (a) Another perspective on the noise, showing significant noise in the enrichment ratio. (b) The beta distribution is used to determine the distribution of fitness values and the Dirichlet distribution is used to determine the distribution of initial populations. We see that these parameters in general have minimal effect on FLIGHTED-Selection log likelihood, but with very low Dirichlet parameters there is a decrease in performance. (c) Left: the $x$-axis is the minimum enrichment ratio and the $y$-axis is the Pearson $r$ between enrichment ratio and fitness for all data points above that minimum. The drastic drop-off in Pearson $r$ as minimum enrichment ratio increases shows that high-activity variants essentially provide $0$ information from just their enrichment ratio. Right: $y$-axis is the number of variants, showing that this is still a significant number of variants. We also tried filtering at a minimum of $10$ reads, which improves results but still shows significant noise.

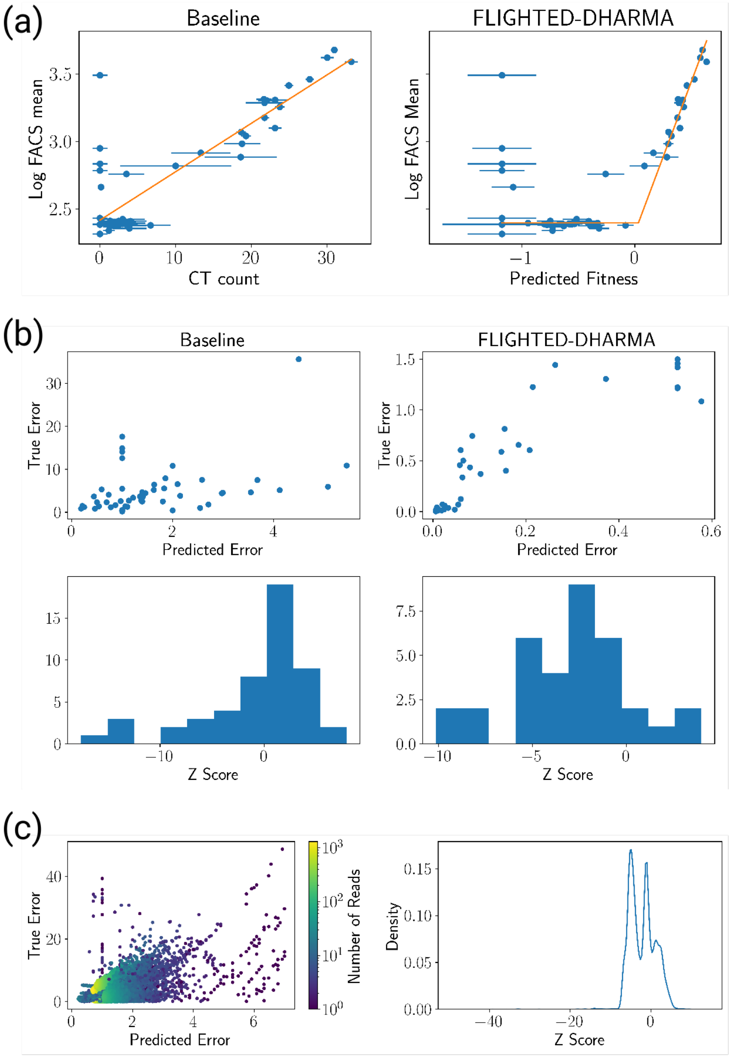


Figure S5: FLIGHTED-DHARMA Model Performance. (a) Accuracy on Validation set for (left) Baseline and (right) FLIGHTED-DHARMA. This shows the fit between the fitness values predicted by each model and the log FACS mean, as done by either a piecewise linear function or a linear function. (b) Calibration of FLIGHTED-DHARMA as Compared to Baseline. The baseline model's true errors are not related to the predicted error, while FLIGHTED-DHARMA has higher true error when predicted error is higher. As such, FLIGHTED-DHARMA is considerably better calibrated with many fewer $z$-score outliers. (c) Calibration of Baseline C$\to$T Model. The baseline model's calibration is considerably worse compared to FLIGHTED-DHARMA's in Figure 3f and g.

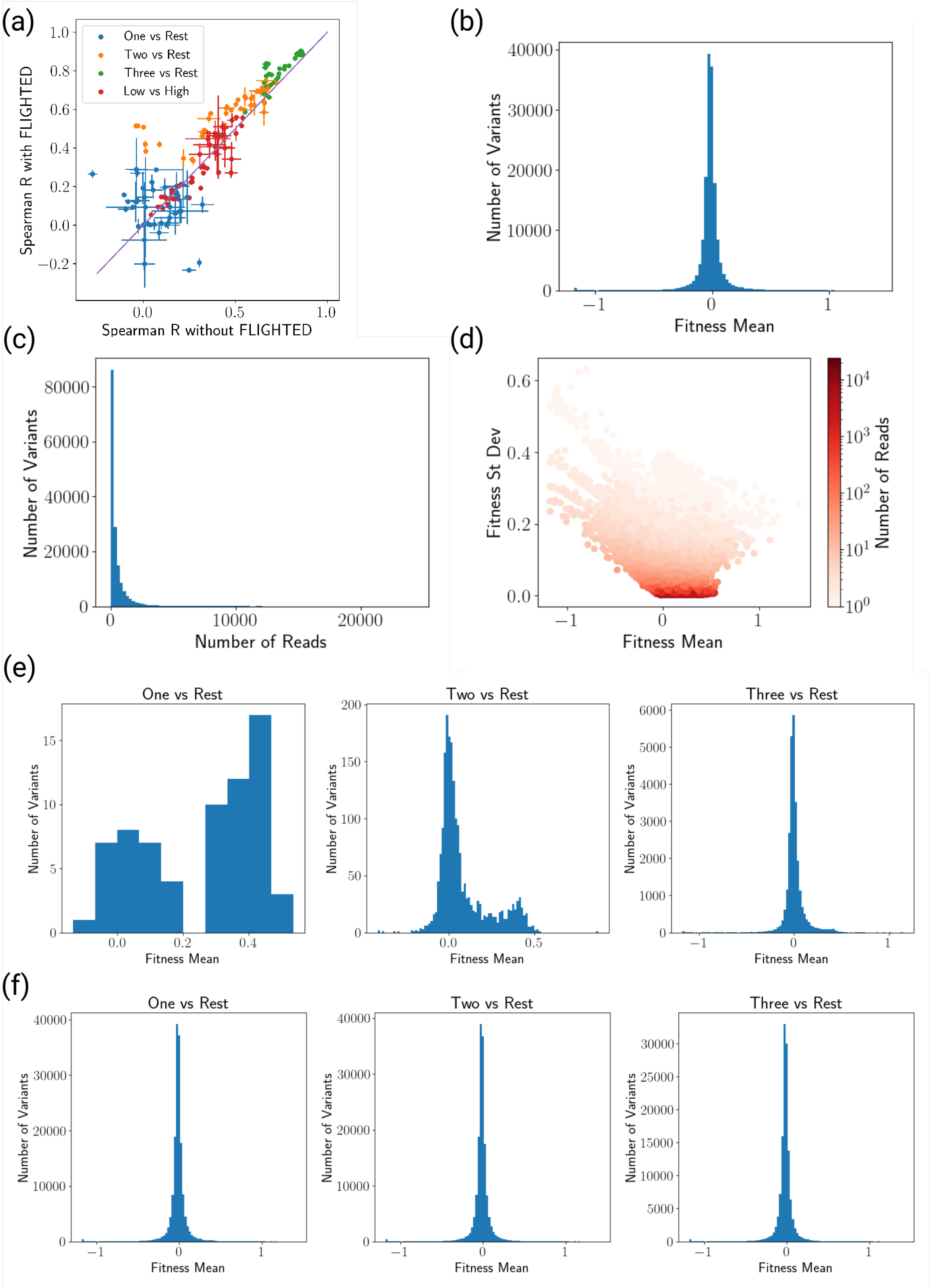


Figure S6: Benchmarking on FLIGHTED Landscapes. (a) FLIGHTED also modestly improves model performance on the GB1 dataset, even when performance is measured as Spearman $\rho$ to the non-FLIGHTED test set. (b) Distribution of fitness values in the TEV dataset. (c) Distribution of read counts in the TEV dataset. (d) Fitness mean, variances, and read count in the TEV dataset. (e) Distribution of fitness values in the training sets for various splits of the TEV dataset. (f) Distribution of fitness values in the test sets for various splits of the TEV dataset.
